# Single Cell Transcriptomics Identifies Distinct Choroid Cell Populations Involved in Visually Guided Eye Growth

**DOI:** 10.1101/2023.05.30.542876

**Authors:** Jody A. Summers, Kenneth L. Jones

## Abstract

Postnatal ocular growth is regulated by a vision-dependent mechanism, termed emmetropization, which acts to minimize refractive error through coordinated growth of the ocular tissues. Many studies suggest that the ocular choroid participates in the emmetropization process via the production of scleral growth regulators that control ocular elongation and refractive development. To elucidate the role of the choroid in emmetropization, we used single-cell RNA sequencing (scRNA-seq) to characterize the cell populations in the chick choroid and compare gene expression changes in these cell populations during conditions in which the eye is undergoing emmetropization. UMAP clustering analysis identified 24 distinct cell clusters in all chick choroids. 7 clusters were identified as fibroblast subpopulations; 5 clusters represented different populations of endothelial cells; 4 clusters were CD45+ macrophages, T cells and B cells; 3 clusters were Schwann cell subpopulations; and 2 clusters were identified as melanocytes. Additionally, single populations of RBCs, plasma cells and neuronal cells were identified. Significant changes in gene expression between control and treated choroids were identified in 17 cell clusters, representing 95% of total choroidal cells. The majority of significant gene expression changes were relatively small (< 2 fold). The highest changes in gene expression were identified in a rare cell population (0.11% - 0.49% of total choroidal cells). This cell population expressed high levels of neuron-specific genes as well as several opsin genes suggestive of a rare neuronal cell population that is potentially light sensitive. Our results, for the first time, provide a comprehensive profile of the major choroidal cell types and their gene expression changes during the process of emmetropization as well as insights into the canonical pathways and upstream regulators that coordinate postnatal ocular growth.

## Introduction

At birth most eyes of diurnal vertebrates are out of focus, likely due to a lack of visual feedback in utero. Postnatally, once the eye begins perceiving retinal images, coordinated growth of the refractive tissues of the eye, the cornea and lens, together with that of the axial length of the eye typically reduces the refractive error of the eye through a process termed, “emmetropization”. Although these basic ideas were first proposed over a century ago (Straub, 1889), the cellular and molecular mechanisms underlying the emmetropization process have remained elusive. Based on clinical and experimental studies in the last 40 years in both animals and humans, we are now beginning elucidate how environmental and behavioral factors stabilize or disrupt ocular growth.

Direct evidence of emmetropization has been provided by numerous animal studies in which modulation of the refractive target with plus and minus lenses results in changes in vitreous chamber depth to align the retinal photoreceptors with the focal plane of the eye (Schaeffel et al., 1988). Additionally, interruption of the emmetropization process as a result of the distortion of visual image quality, either through ocular pathology in humans (O’Leary and Millodot, 1979, Rabin et al., 1981, Rasooly and BenEzra, 1988, Twomey et al., 1990) or application of translucent occluders in animal models (Wallman et al., 1978), results in axial elongation and the development of myopia. Form deprivation-induced myopia is reversible; restoration of unrestricted vision (and the resultant myopic defocus) results in a temporary cessation of axial growth, eventually leading to the reestablishment of emmetropia (recovery) in the formerly deprived eye (Wallman and Adams, 1987). The response to deprivation and defocus is rapid, leading to detectable changes in vitreous chamber depth within hours (Zhu et al., 2005). It has been well established that in chicks these visually induced changes in ocular growth are directly associated with changes in proteoglycan synthesis and proteoglycan accumulation in the sclera at the posterior pole of the eye (Christensen and Wallman, 1991; Gentle et al., 2001; McBrien et al., 1991; Rada et al., 1991; 2002).

Although the chicken eye has several significant anatomical differences from eyes of placental mammals (most notably a lack of retinal circulation and the presence of a cartilaginous layer in the sclera), virtually all of the important observations on emmetropization initially made in chicks have been reproduced in higher animals. These findings include: 1) The mechanisms underlying emmetropization are found locally, within the eye, and that emmetropization can occur in the absence of an intact optic nerve (McFadden and Wildsoet, 2020; Norton et al., 1994; Troilo et al., 1987); 2) Eyes can compensate both for imposed positive (myopic) and negative (hyperopic) defocus (Schaeffel et al., 1988; Troilo et al., 2009); 3) Form deprivation restricted to localized regions of the retina can stimulate scleral remodeling in regions adjacent to the region of deprived retina (Smith et al., 2009; Wallman, Gottlieb et al., 1987); 4) Choroidal thickening occurs during compensation of myopic defocus, either during recovery from induced myopia or following application of positive lenses to normal eyes (Troilo et al., 2000; Wallman et al., 1995); and 5) Changes in scleral extracellular matrix (ECM) synthesis and remodeling are responsible for changes in eye length (Norton and Rada, 1995; Rada et al., 1991, 2000). From these and many other studies, it is now generally accepted that emmetropization is regulated by a retina-to-scleral chemical cascade that transmits signals, initiated in the retina in response to visual stimuli, to the sclera to effect changes scleral ECM remodeling, axial length and refraction (Summers, Schaeffel et al., 2021).

Of much interest are the choroidal changes associated with emmetropization, as any retinal-derived scleral growth regulator must pass through the choroid, or act on the choroid to synthesize additional molecular signals that can subsequently act on the sclera to stimulate scleral ECM remodeling. Several studies have characterized the choroidal changes associated with recovery and compensation for myopic defocus. Choroidal thickening (Wallman et al., 1995), increased choroidal permeability (Pendrak et al., 2000; Rada and Palmer, 2007), and increased choroidal blood flow (Fitzgerald et al., 2002; Jin and Stjernschantz, 2000) have been well-documented during recovery from induced myopia. Additionally, increased choroidal synthesis of retinoic acid (Mertz and Wallman, 2000; Troilo et al., 2006), the retinoic-acid synthesizing enzyme, ALDH1a2 (Harper et al., 2016), ovotransferrin (Rada et al., 2001), and apolipoprotein A-I (Summers et al., 2016) have been observed during recovery or compensation to positive lenses. The choroid is a complex tissue, consisting of a rich blood supply, lymphatic vessels, stromal cells, intrinsic choroidal neurons, extravascular smooth muscle and axons of sympathetic, parasympathetic and sensory neurons (Nickla and Wallman, 2010). Therefore, to gain deeper insight into the choroidal response that accompanies recovery and compensation for myopic defocus, the present study was undertaken to: 1) identify the major choroidal cell clusters in control chicken eyes and eyes recovering from induced myopia, and 2) Investigate the gene expression profiles of each choroidal cell population and how they change in response to myopic defocus/recovery. Our data, for the first time, identify specific choroidal cell types that undergo transcriptional changes in response to myopic defocus/recovery and provide us with additional insights into the key mediators in the retina-to-sclera chemical cascade during the regulation of eye growth.

## Results

### Cell Populations Identified in Chick Choroids

To investigate the gene expression profiles of chick choroid cell populations, we dispersed the cells of choroids from control and recovering eyes of 15 day old male chicks, isolated live cells from dead cells, and performed single-cell RNA-seq (scRNA-Seq) analysis using the 10x Genomics system (Macosko et al., 2015) **(Figure 1)**. FACS analyses indicated that 89 – 94 % of cells in each sample were living at the time of isolation, based on their incorporation of calcein (green fluorescence) and were sorted at final concentrations ranging from 1.11 x 10^6^ – 8.75 x 10^6^ cells/ml (**Figure 2 and Figure 2 – figure supplement 1**).

**Figure 1.**
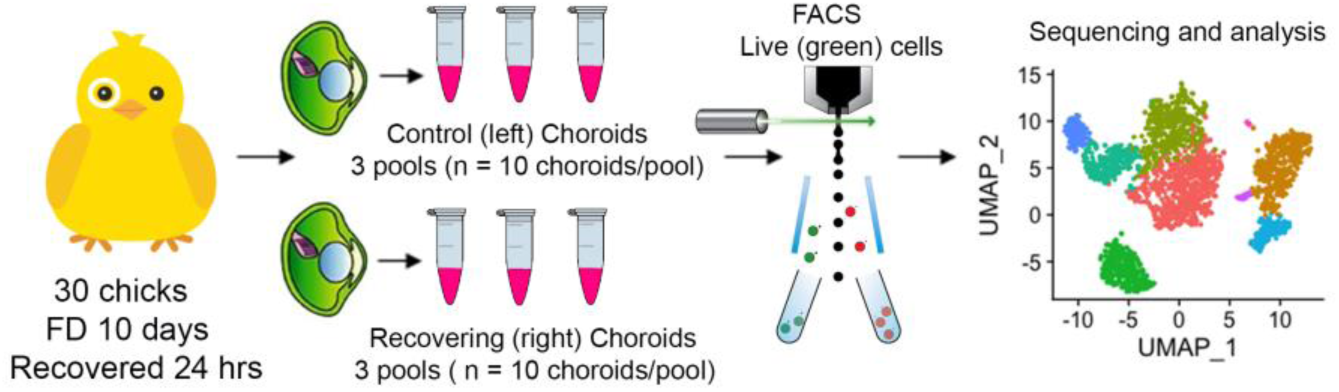
Single-cell analysis of chick choroids. Choroids were isolated from control (left) eyes and eyes recovering from form deprivation myopia (right) and pooled into six groups of 10 choroids each (3 control pools and 3 recovering pools). Choroids were enzymatically and mechanically dispersed, living cells (labeled with calcein AM) were isolated using fluorescence activated cell sorting (FACS), and subjected to single cell RNA sequencing and bioinformatic analysis.

**Figure 2.**
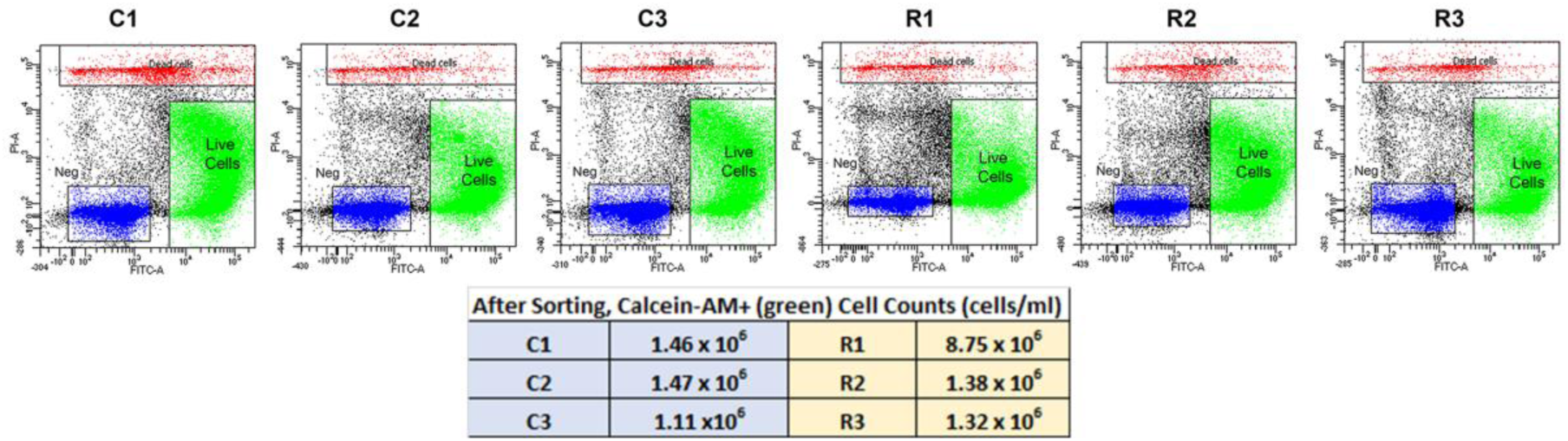
FACS Isolation of Choroidal Cells from Control (C1,C2 and C3) and Recovering (R1, R2 and R3) Eyes. Living choroidal cells were purified via fluorescence-activated cell sorting (FACS) based on single-cell populations identified by forward scatter (FSC), single-cell populations by side scatter (SSC), cells fluorescently labeled with calcein AM (FITC-A; green), and not labelled with EthDIII (PI-A; red).

A total of 71,054 choroid cells passed the quality control check: 10,221, 11,251 and 12,469 cells for samples C1, C2 and C3, respectively (three choroids/sample) and 11,976, 12,392 and 12,745 cells for samples R1, R2 and R3, respectively (three choroids/sample) (**Table S1**).

### Marker Genes Identified in Chick Choroids

A UMAP plot consisting of 24 distinct cell clusters was developed in Seurat based on similarity of gene expression and statistically significant gene expression differences within and between cell clusters using PCA/UMAP in Seurat. A biomarker list for each cluster was generated using T-test in Seurat (FDR < 0.05; **Table 1 and Table 1 – figure supplement 1**). Using the top differentially expressed genes from each cluster and well-established cell-specific markers, we identified and annotated 24 clusters of cells in the chick choroid (**Table 2, Figure 3**).

**Figure 3.**
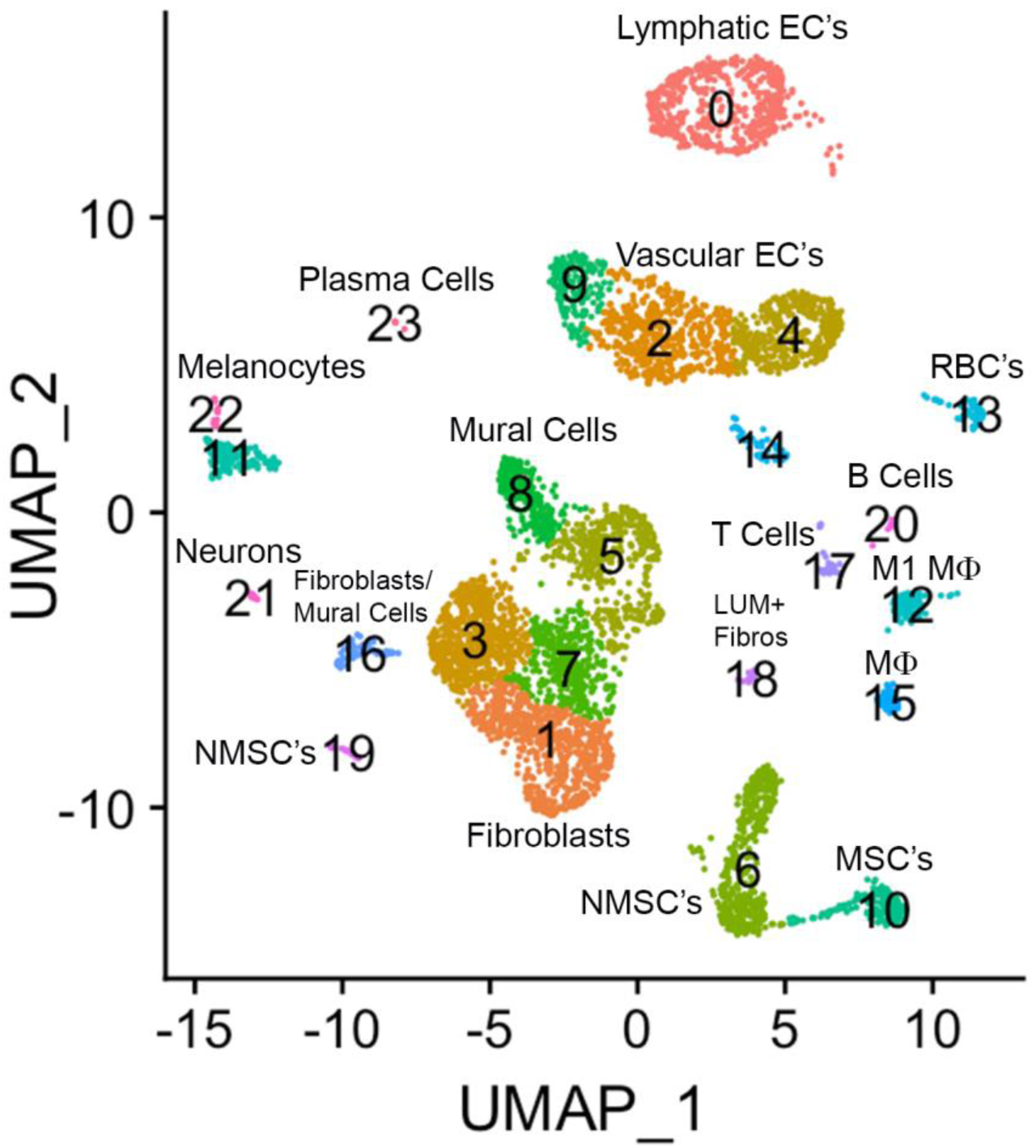
The major cell clusters in the chicken choroid. Uniform Manifold Approximation and Projection (UMAP) map showing the identified 24 distinct choroid cell types based on the transcriptomes of 71,054 cells. Cells are colored by Seurat clustering and annotated by cell types (each point represents a single cell). EC’s, endothelial cells; MSC’s, myelinating Schwann cells; NMSC’s, non-myelinating Schwann cells; LUM+ fibros, lumican-positive fibroblasts; Mɸ, macrophage; RBC’s, red blood cells.

**TABLE 1.** Selected marker genes identified in 24 chicken choroid cell clusters.

| **Cell cluster** | **Selected marker genes** |
| --- | --- |
| 0 | *CPE, MMRN1, PROX1, LYVE1, GALNT1* |
| 1 | *MGP, CYTL1, TCF15ENS, ANGPT2, VIM* |
| 2 | *SOX17, DIO2, SMS, TCF15, SLC6A6* |
| 3 | *PTGDS, IL1RL1, CHGA, COL8A1, NCAM1* |
| 4 | *CA4, ENSGALG0000007646, EHD3, RGS2, SIX3* |
| 5 | *MYH11, CLEC3B, RGS5, ACTA2, MYLK* |
| 6 | *PRNP, CDH19, CRYAB, SLIT1, ENSGALG00000010722* |
| 7 | *FGFR3, BIRC3, TAGLN, COL1A1* |
| 8 | *RGS4, RERGL, KCNJ8, HTRA1, ENSGALG00000031430* |
| 9 | *VWE, NPR3, FILIP1, VEGFC, MBP* |
| 10 | *ENSGALG00000047143, PMP22, MPZ, FABP9, CRP* |
| 11 | *MLANA, PMEL, GPNMB, TYRP1, IL1B* |
| 12 | *AVD, CCL4, ENSGALG0000002431, ENSGALG00000053121, HBBA* |
| 13 | *HBA1, HBAD, IFI27L2, HBE1, SMC2* |
| 14 | *CENPF, ENSGALG00000025996, CENPE, ASPM, CD74* |
| 15 | *IL8L1, BLB2, BLB1, TMSB15B, TOP2A* |
| 16 | *CENPF, ENSGALG00000025996, CENPE, ASPM, CD74* |
| 17 | *CD3D, ENSGALG00000041611, XCL1, PTPRC, LUM* |
| 18 | *ENSGALG00000035854, TNMD, EBF2, SCARA5, CKAP2* |
| 19 | *CENPF, ENSGALG00000025996, CENPE, ASPM, CD74* |
| 20 | *JCHAIN, ENSGALG00000011190, ZP1, ENSGALG00000046212, LPAR5* |
| 21 | *GAL, CHRNA3, SCN9A, STMN2, EEF1A2* |
| 22 | *EDNRB2, TYR, TUBB3, PRLH* |
| 23 | *ENSGALG00000049450, ENSGALG00000049716, SH2D6, POU2AF1* |

**Table 2.**
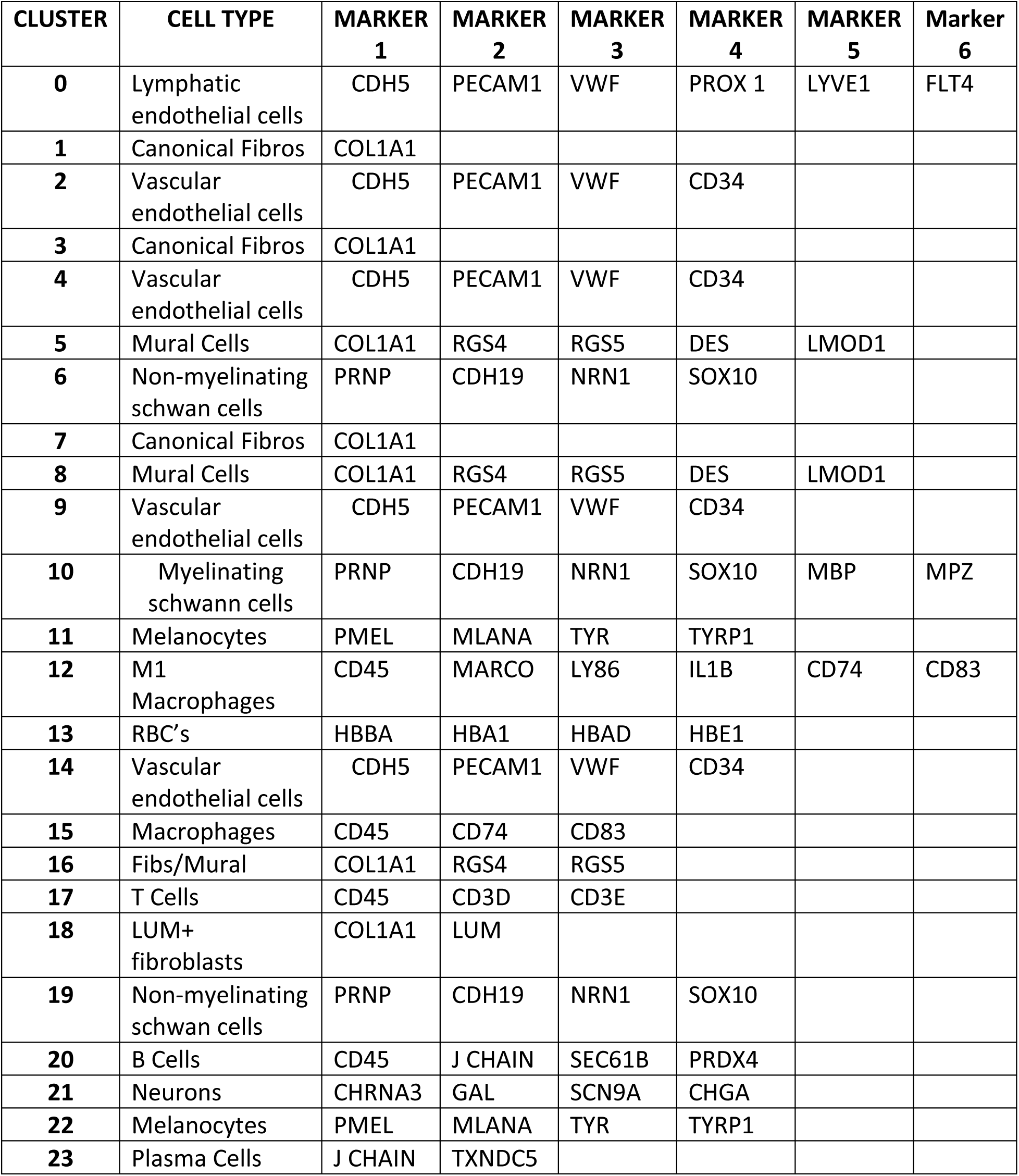
Identification of Major Choroidal Cell Types using Established Marker Genes

| **CLUSTER** | **CELL TYPE** | **MARKER**<br>**1** | **MARKER**<br>**2** | **MARKER**<br>**3** | **MARKER**<br>**4** | **MARKER**<br>**5** | **Marker**<br>**6** |
| --- | --- | --- | --- | --- | --- | --- | --- |
| **0** | Lymphatic endothelial cells | CDH5 | PECAM1 | VWF | PROX 1 | LYVE1 | FLT4 |
| **1** | Canonical Fibros | COL1A1 |  |  |  |  |  |
| **2** | Vascular endothelial cells | CDH5 | PECAM1 | VWF | CD34 |  |  |
| **3** | Canonical Fibros | COL1A1 |  |  |  |  |  |
| **4** | Vascular endothelial cells | CDH5 | PECAM1 | VWF | CD34 |  |  |
| **5** | Mural Cells | COL1A1 | RGS4 | RGS5 | DES | LMOD1 |  |
| **6** | Non-myelinating schwann cells | PRNP | CDH19 | NRN1 | SOX10 |  |  |
| **7** | Canonical Fibros | COL1A1 |  |  |  |  |  |
| **8** | Mural Cells | COL1A1 | RGS4 | RGS5 | DES | LMOD1 |  |
| **9** | Vascular endothelial cells | CDH5 | PECAM1 | VWF | CD34 |  |  |
| **10** | Myelinating schwann cells | PRNP | CDH19 | NRN1 | SOX10 | MBP | MPZ |
| **11** | Melanocytes | PMEL | MLANA | TYR | TYRP1 |  |  |
| **12** | M1 Macrophages | CD45 | MARCO | LY86 | IL1B | CD74 | CD83 |
| **13** | RBC's | HBBA | HBA1 | HBAD | HBE1 |  |  |
| **14** | Vascular endothelial cells | CDH5 | PECAM1 | VWF | CD34 |  |  |
| **15** | Macrophages | CD45 | CD74 | CD83 |  |  |  |
| **16** | Fibs/Mural | COL1A1 | RGS4 | RGS5 |  |  |  |
| **17** | T Cells | CD45 | CD3D | CD3E |  |  |  |
| **18** | LUM+ fibroblasts | COL1A1 | LUM |  |  |  |  |
| **19** | Non-myelinating schwann cells | PRNP | CDH19 | NRN1 | SOX10 |  |  |
| **20** | B Cells | CD45 | J CHAIN | SEC61B | PRDX4 |  |  |
| **21** | Neurons | CHRNA3 | GAL | SCN9A | CHGA |  |  |
| **22** | Melanocytes | PMEL | MLANA | TYR | TYRP1 |  |  |
| **23** | Plasma Cells | J CHAIN | TXNDC5 |  |  |  |  |

Clusters 0, 2, 4, 9, and 14 were determined to be different populations of endothelial cells, based on expression of several endothelial cell (EC) markers (CDH5, PECAM1 and VWF). Additionally, Cluster 0 expressed markers for Lymphatic ECs (PROX1, LYVE1 and FLT4), while clusters 9, 2, 4, and 14 expressed markers for classical vascular ECs. The continuum of clusters 9 → 2 → 4 represent ECs from vessels of differing vascular size, where Cluster 9 is represents EC from large vessels, and Cluster 4 are capillary ECs based on the relative expression of the markers PECAM1, VWF and CD34. PECAM1 and VWF, markers known to be expressed in ECs of larger blood vessels, were highly expressed in cluster 9, and had the lowest expression in cluster 4; while CD34, a marker generally associated with ECs of capillaries and nascent ECs, was expressed at the highest level in Cluster 4 (Pusztaszeri et al., 2006). Clusters 22 and 11 were identified as melanocytes; cluster 22 cells are melanocytes that are exclusively in the cell cycle phases G2/M, whereas cluster 11 melanocytes in cell cycle phases G1 and S (**Figure 3– figure supplement 1**). Cluster 13 represented red blood cells based on expression of several hemoglobin subunits (HBBA: hemoglobin subunit epsilon 1; HBA1: hemoglobin subunit alpha 1; HBAD: hemoglobin alpha, subunit D; and HBE1: hemoglobin subunit epsilon 1). Clusters 6, 10 and 19 were identified as Schwann Cells based on the expression of PRNP, CDH19, NRN1 and SOX10. Additionally, cells in Cluster 10 also express the myelin genes, MBP (myelin basic protein) and MPZ (myelin protein zero), indicating that cluster 10 cells are myelinating Schwann cells while clusters 6 and 19 represent non-myelinating Schwann cells. Cluster 19 represents non-myelinating Schwann cells in cell cycle stage G2M, based on expression of a number of G2M and S phase gene markers in the “Cell Cycle Scoring” function (Seurat 4.0.6) while non-myelinating Schwann cells in cluster 6 represent cells in stages G1 and S of the cell cycle (**Figure 3 – Figure supplement 1**).

Clusters 12 and 15 were identified as macrophages/dendritic cells, based on expression of CD45, CD74 and CD83. Cluster 12 cells represent M1 macrophages as they express the macrophage markers MARCO and LY86 as well as the pro-inflammatory cytokine, IL1B. We believe that cluster 15 cells are M2 macrophages, however the chicken genome that was used for the analysis has a limited number of markers for M2 macrophages. Therefore, we chose to refer to cluster 15 cells as “Macrophages, not M1”. Cluster 17 cells were identified as T lymphocytes (T cells) based on their expression of CD45, CD3D and CD3E, while Cluster 20 cells are B lymphocytes (B cells), based on their expression of CD45, J CHAIN, SEC61B and PRDX4. Cluster 23 represents plasma cells based on relatively high expression of J CHAIN and TXNDC5. Cluster 21 cells were determined to be neurons/neuroendocrine cells based on their expression of CHRNA3, GAL, SCN9A, and CHGA.

Clusters 8,5,3,7,1,16,18 were determined to be fibroblast-like cells expressing COL1A1. Clusters 3,7,1 are canonical fibroblasts, while clusters 8 and 5 are mural cells (smooth muscle cells and pericytes) based on their expression of RGS4, RGS5, DES, LMOD1, in addition to COL1A1 (Betsholtz 2020; Muhl et al., 2020). Cluster 18 was identified as a subpopulation of fibroblasts that express the proteoglycan, lumican (LUM). Cluster 16 appears to be a combination of mural and canonical fibroblasts that are in an alternate cell cycle (**Figure 3– figure supplement 1**). By comparing RGS4/5+ cells with that of COL1A1+ cells it can be observed that mural cells are located in the upper region of cluster 16, whereas canonical fibroblasts are located in the bottom region of cluster 16 (**Figure 3 – figure supplement 2**).

The number of clusters and number of cells per cluster were similar for most cell clusters between the three control and three recovering samples (**Figure 4A**), however 5 cell clusters displayed statistically significant differences in the percentage of total cells between control and recovering samples (**Figure 4B, 4C**). Significant increases in cell numbers were noted in recovering samples among lymphatic endothelial cells (cluster 0), vascular endothelial cells (clusters 9 and 14) and M1 macrophages (cluster 12). Additionally, a significant decrease in the percentage of non-myelinating Schwann cells was observed in recovering samples.

**Figure 4.**
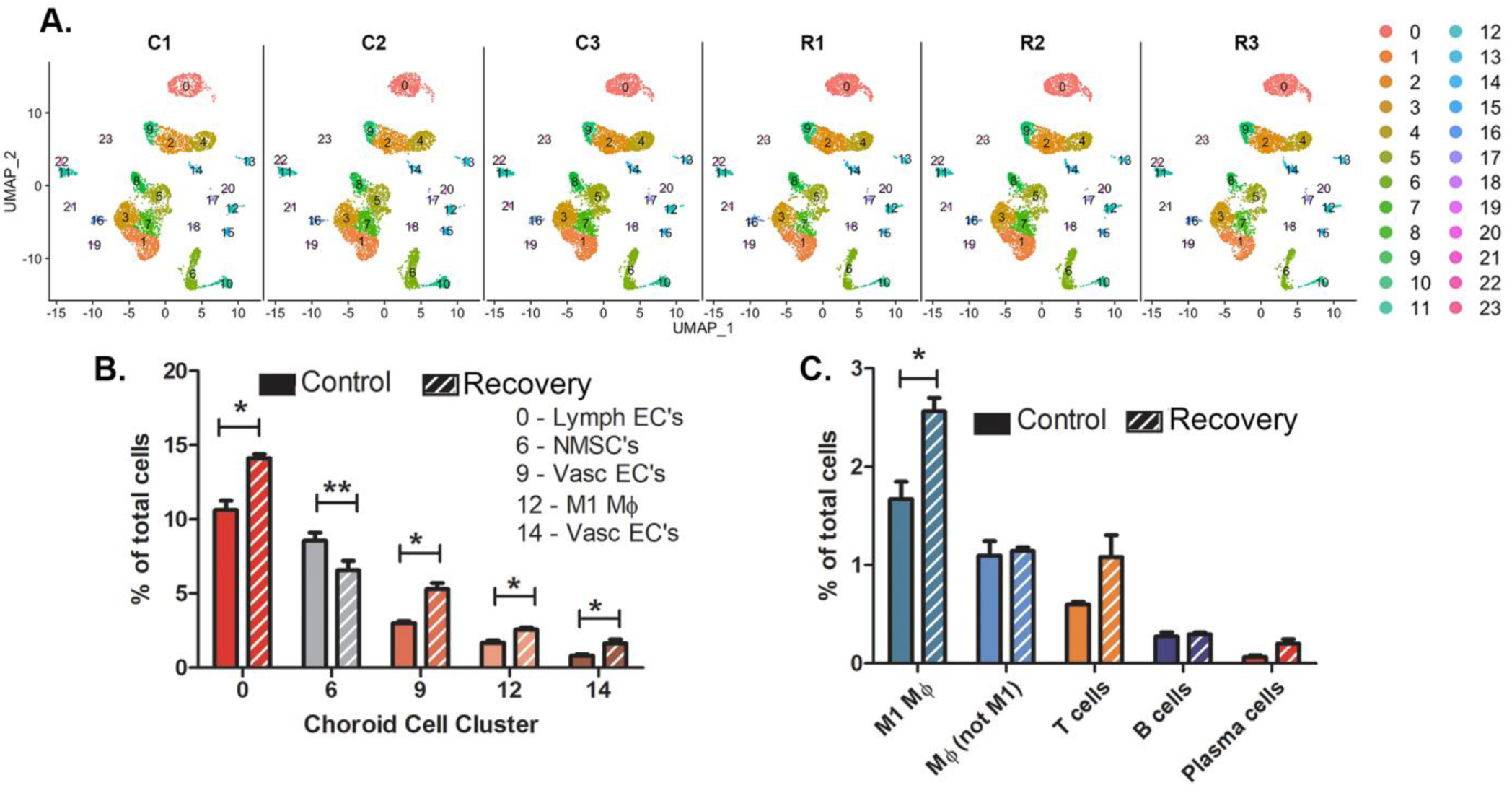
There are limited differences in choroid cell clusters between control and recovering chick eyes. **A)** Uniform Manifold Approximation and Projection (UMAP) of 24 choroid cell clusters (0 – 23) from three pools of control eyes (C1, C2, C3; 10 choroids/pool) and from three pools of recovering eyes (R1, R2, R3; 10 choroids/pool). **B)**. Comparison of the number of cells in clusters between control and recovering choroids (only clusters with significant differences in cell numbers are shown). **C)** Comparison of immune cell types between control and recovering choroids. **p < 0.01; *p < 0.05, t-test. Data are presented as percentages of the total cells analyzed in each sample.

Additionally, we compared the numbers of immune cells in control and recovering choroid samples (as a percentage of the total number of cells/sample) (**Figure 4C**). As previously noted, we observed a significant increase in the number of M1 macrophages in recovering choroids as compared with control choroids (p < 0.05, paired t-test). This M1 macrophage population was the largest of all immune cell populations in both control and recovering samples. No significant differences were detected in the other macrophage population (cluster 15, Mɸ, not M1), T cells, B cells or plasmas cells between recovering and control choroid samples (as a percentage of the total number of cells/sample).

### Gene Expression Differences Associated with Recovery from Induced Myopia

Gene expression [expressed in fragments per kilobase of exon per million mapped fragments (FPKM)] in all 24 choroid cell clusters was compared between the three control and three recovering choroid samples. Significant changes in gene expression were observed in 17 of the 24 cell clusters (**Figure 5**). No significant differences in gene expression were detected in RBC’s (cluster 13), macrophages, not M1 (cluster 15), T cells (cluster 17), non-myelinating Schwann cells in cell cycle stage G2M (cluster 19), B cells (cluster 20), melanocytes in the cell cycle phases G2/M (cluster 22) and plasma cells (cluster 23). However, due to the large differences in cell numbers within each choroid cell population (e.g. population 0 is the largest population with 513 – 949 cells and population 23 is the smallest with 2 – 20 cells), statistical significance could be reached with smaller differences in gene expression among clusters with larger sample sizes and only genes with large fold changes could be reported as significantly different in clusters with smaller sample sizes. Relatively small, but significant, fold changes (< 2.5 fold) were detected in many genes between control and recovering choroidal cell clusters. Greater fold changes were observed for NOV (CCN3) in cluster 3 and cluster 16; AVD (avidin) in cluster 12; DIO2 (Type II iodothyronine deiodinase) in cluster 14; RGS16 (regulator of G protein signaling 16) and NPR3.00 (natriuretic peptide receptor 3) in cluster 16; and four of the five genes differentially expressed in cluster 21 [SLC38A2 (sodium-coupled neutral amino acid transporter 2), CST3 (cystatin c) CTSV (cathepsin V) and ENSGALG00000017040 (complement C4A)].

**Figure 5.**
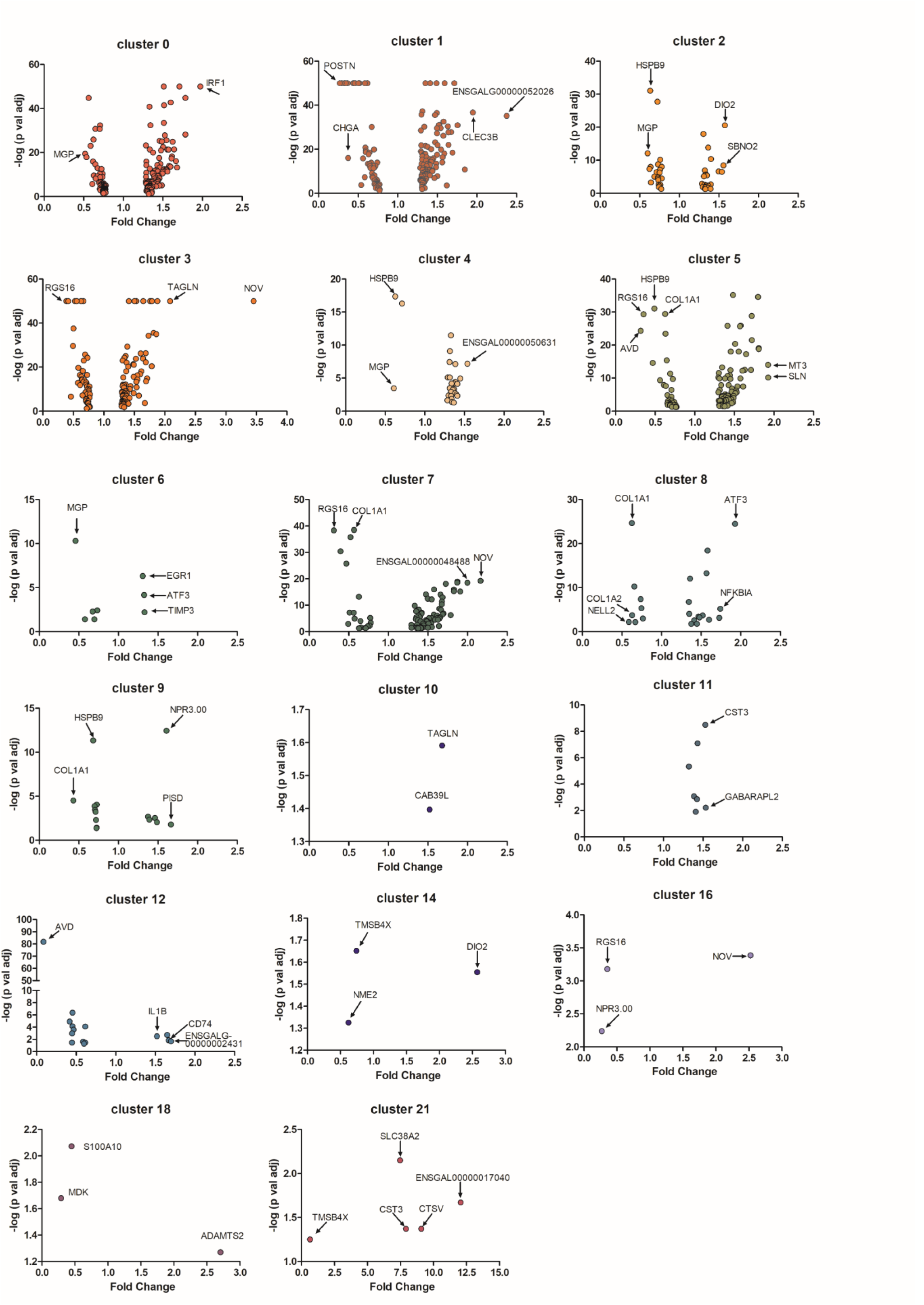
Differential gene expression in choroid cell clusters from control and recovering eyes. Volcano plots demonstrate fold changes and p values of individual genes in 17 choroidal cell clusters. Only differentially expressed genes with a p-value of ≤ 0.05 are represented (t-test, n = 3). Genes with relatively large fold changes and/or very small p values are labelled and indicated with arrows. MGP, matrix gla protein; IRF1, interferon regulatory factor 1; POSTN, periostin; CHGA, chromogranin A; CLEC3B, c-type lectin domain family 3 member B; ENSGALG00000052026, novel gene, matched to serine peptidase inhibitor, Kazal type 2; HSPB9, heat shock protein family B (small) sember 9; DIO2, iodothyronine deiodinase 2; SBNO2, strawberry notch homolog 2; RGS16, regulator of g protein signaling 16; TAGLN, transgelin; NOV, nephroblastoma overexpressed; ENSGALG00000050631, novel gene; AVD, avidin; COL1A1, collagen type I alpha 1 chain; MT3, metallothionein 3; SLN, sarcolipin; EGR1, early growth response 1; ATF3, activating transcription factor 3 ; TIMP3, TIMP metallopeptidase inhibitor 3; ENSGALG00000048488, novel gene; COL1A2, collagen type I alpha 2 chain; NELL2, neural EGFL like 2; NFKB1A, nuclear factor kappa B subunit 1; NPR3.00, natriuretic peptide receptor 3; PISD, phosphatidylserine decarboxylase; CAB39L, calcium binding protein 39 like; CST3, cystatin c; GABARAPL2, GABA type A receptor associated protein like 2; IL1B, interleukin 1 beta; CD74, cluster of differentiation 74; ENSGALG00000002431, novel gene, matched to complement factor H; TMSB4X, thymosin beta 4 X-Linked; NME2, nucleoside diphosphate kinase B; S100A10, S100 calcium binding protein A10; MDK, midkine; ADAMTS2, ADAM metallopeptidase with thrombospondin type 1 motif 2; SLC38A2, solute carrier family 38 member 2; CTSV, cathepsin V; ENSGALG00000017040, novel gene, matched to complement C4A.

In order to obtain an overview of the differentially expressed genes in choroidal cell clusters, a heat map was generated for all genes that were significantly up or down regulated in three or more cell clusters (**Figure 6**). This heatmap demonstrated that for any particular gene, recovery-induced changes in gene expression were similar across multiple cell clusters; i.e. genes that were up-regulated in one cell cluster were upregulated in multiple cell clusters and genes that were downregulated were downregulated in multiple cell clusters. Only 4 genes showed upregulation in some clusters and downregulation in others (CYR61, ACTB, CD74 and NPR3.00). Also evident from this heatmap are the highly overexpressed genes in cluster 21, CST3 and CTSV, which showed the highest fold increases of all differentially expressed genes, indicated in red (the other two highly expressed genes in cluster 21, SLC38A2 and ENSGALG00000017040, were not expressed in at least 3 choroidal cell clusters and therefore not included in the heatmap). The genes, EGR1 and ATF3 were the most globally upregulated genes, found to be upregulated in 8 and 7 individual choroid cell clusters, respectively. Genes COL1A1 and HSPB9 were the most globally downregulated genes, both found to be downregulated in 9 individual cell clusters.

**Figure 6.**
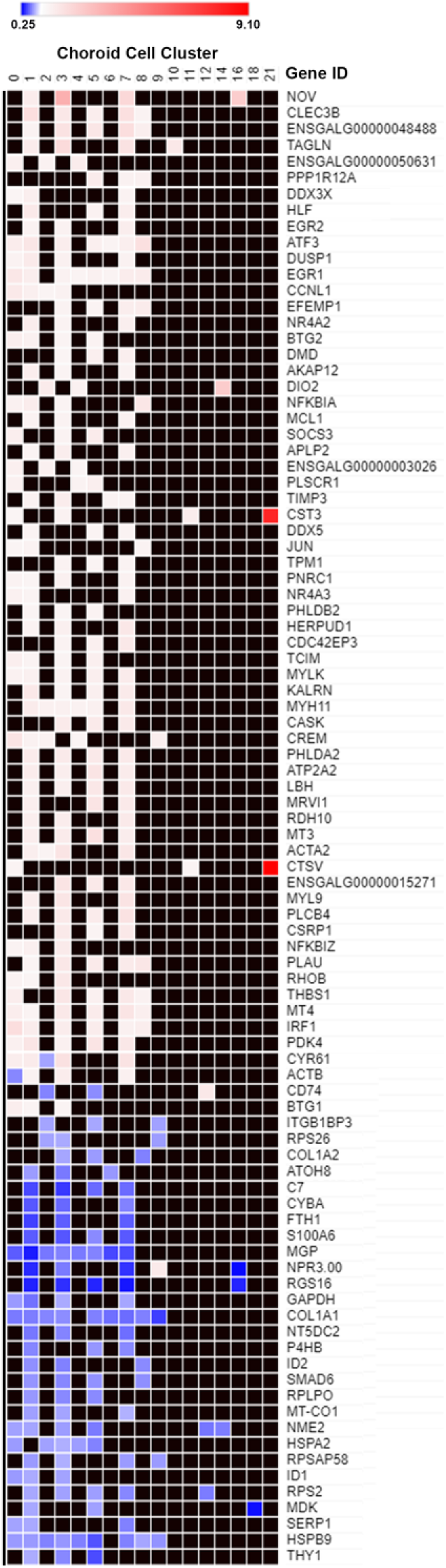
Differentially expressed genes in multiple choroid cell clusters. Genes that were significantly up or down-regulated in 3 or more choroid cell clusters (data from Figure 5) were plotted on a heat map. The blue shades denote genes down-regulated in recovering choroids compared with controls and white, pink and red shades denote genes up-regulated in recovering choroids compared with controls. Black squares indicate no differential expression.

### Cluster 21 (Neurons)

Although cluster 21 represented a rare cell population (7 – 35 cells/ cluster), the large fold changes in expression of several genes by these cells in recovering choroids prompted us to investigate this population of cells further. **Figure 7** shows several genes enriched by this population, compared with their expression in all other clusters. As expected, many genes enriched in cluster 21 are associated with neuronal cells and were used to identify cluster 21 cells as neurons (**Tables 1, 2**).

**Figure 7.**
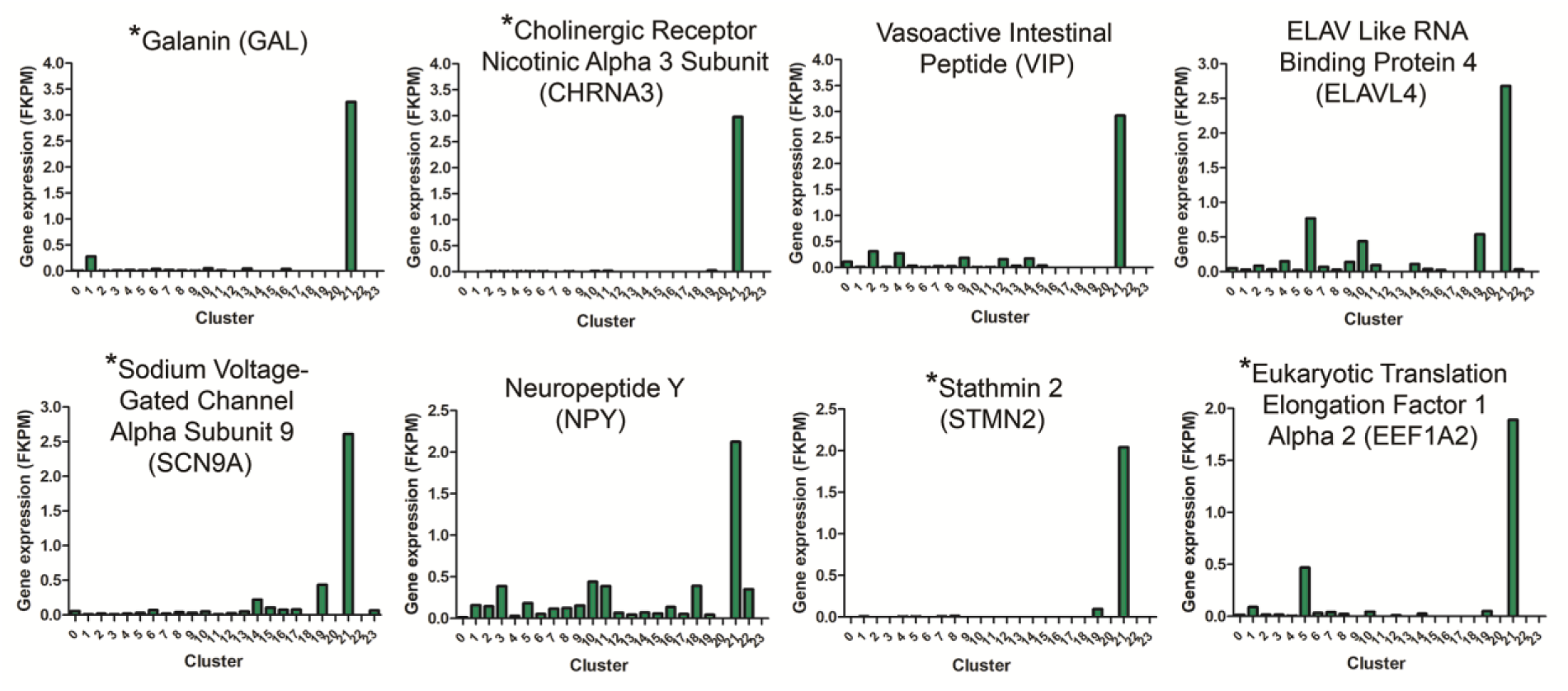
Gene expression signatures for cluster 21 neurons. Genes upregulated in cluster 21 relative to all other clusters are indicated based on their FKPM values. Gene names indicated with an asterisk (*) are genes used to distinguish cluster 21 from other choroidal cell clusters (Table 2).

We were able to identify these cells in choroidal whole mounts, following immunolabelling with anti-CHRNA3. CHRNA3-positive cells were very sparsely distributed throughout the choroidal stroma and appeared as small round cells or cells with one or more elongated processes in choroids from control and recovering eyes (**Figure 8A**). CHRNA3-positive cells were also isolated from cell suspensions of choroid tissue digests of normal chicken eyes, following fixation and immunolabelling with anti-CHRNA3, using fluorescence activated cell sorting (FACS). Quantification of the CHRNA3-positive cells indicated that CHRNA3-positive cells represented ≈ 0.49% of all choroidal cells [CHRNA3/Alexa 647/ DAPI labelled cells (0.60%) minus non-specifically labelled cells (0.11%)] (**Figure 8b**), which is close to the percentage of cluster 21 cells in choroidal samples of control and recovering eyes determined by scRNA seq (0.11 - 0.48 %, **supplementary Table S2**).

**Figure 8.**
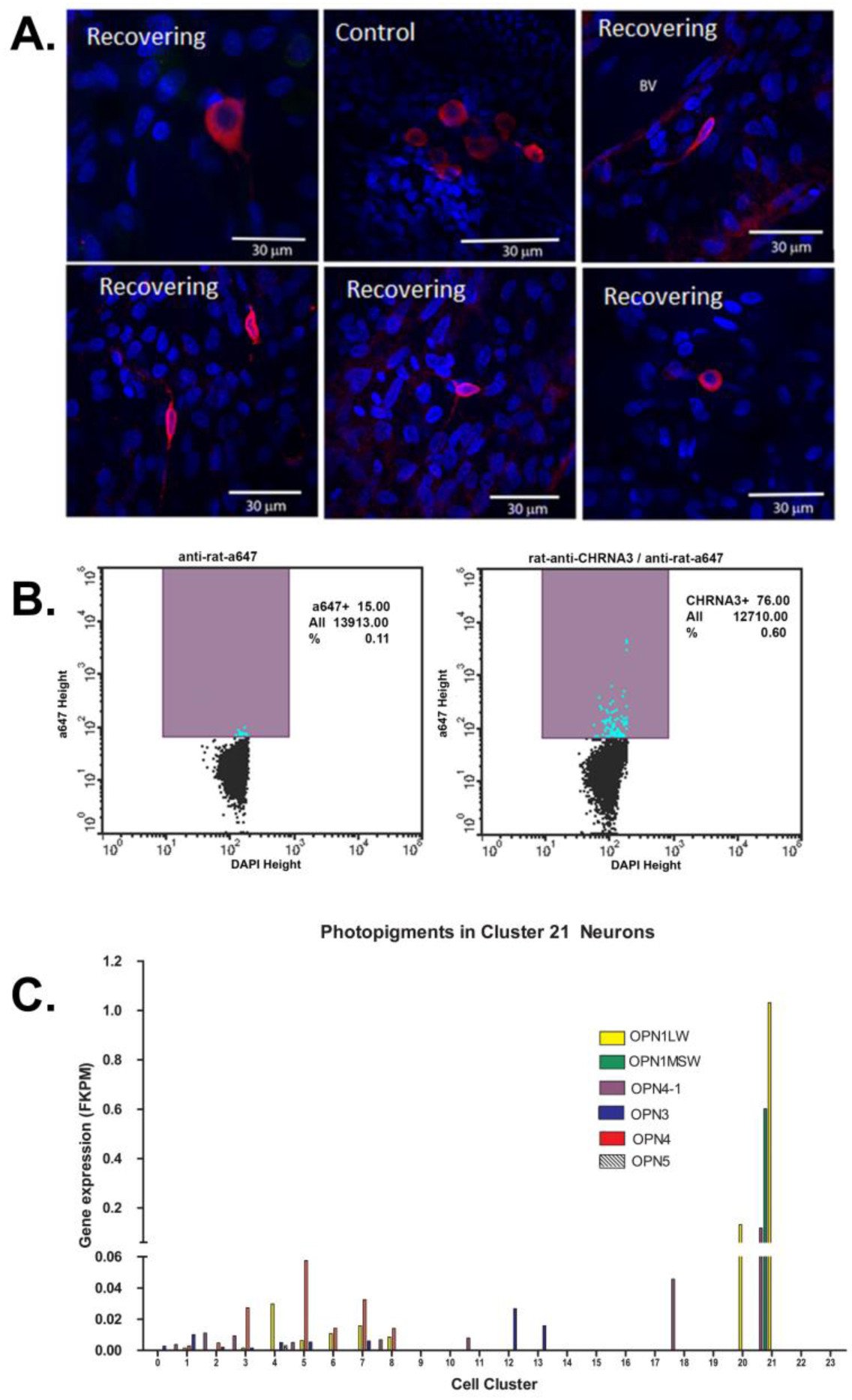
Cluster 21 cells are a rare population of opsin-expressing neurons. **A.** Anti-CHRNA3 antibodies were used to identify cluster 21 neurons in whole mounts of chick choroids from control and recovering eyes. CHRNA3-positive cells were labeled with anti-rat AlexaFluor 568 (red). Nuclei are stained with DAPI (blue). Scale bars are 30 μm. **B.** CHRNA3-positive cells were isolated from choroid tissue homogenates from normal chicken eyes by FACS, following labelling with anti-CHRNA3 and anti-rat-AlexaFluor 647 and DAPI staining of nuclei. CHRNA3/AlexaFluor 647/DAPI positive cells represented 0.60% of the total DAPI-positive cells. Non-specifically labeled cells, in which cells were labelled only with AlexaFluor 647 and DAPI represented 0.11% of the total DAPI-positive cells. **C.** Opsin gene expression (FKPM) was compared among the 24 choroid cell clusters. OPN1LW, opsin 1, long wave sensitive; OPN1MSW, opsin, green sensitive; OPN4-1, photopigment, melanopsin-like; OPN3, opsin 3; OPN4, opsin 4, melanopsin; OPN4, opsin 5.

It has been suggested that the choroid may be intrinsically light sensitive since the choroid has been shown to undergo thickness changes in response to different wavelengths of light (Lou and Ostrin, 2020; Lou and Ostrin, 2023). We therefore evaluated the expression of light sensitive photopigments (opsin) genes in all cell clusters in the chick choroid (**Figure 8c**). The opsin genes, OPN3, OPN4, OPN4-1, OPN5, OPN1MSW and OPN1LW were expressed in one or more cell clusters in the chick choroid. Interestingly, opsins OPN4-1, OPN1MSW and OPN1LW were highly overexpressed in Cluster 21 as compared with their expression in other cell clusters, and ON1MSW was exclusively expressed in Cluster 21. Taken together, these data suggest that cluster 21 cells may represent a population of light sensitive neurons in the choroid.

### Identification of Canonical Pathways and Upstream Regulators

Ingenuity Pathway Analyses was used to identify common canonical pathways involved in the choroidal recovery response and common upstream regulators that may mediate the visually-induced gene expression differences within each choroidal cell cluster between control and recovering eyes. Clusters with less than six significantly differentially expressed genes could not be included in the analyses. Therefore, comparison analyses of canonical pathways and upstream regulators were carried out for 12 cell clusters (clusters 0, 1, 2, 3, 4, 5, 6, 7, 8, 9, 11, and 12) and heat maps were generated and visualized by z-scores and sorted by hierarchical clustering **(Figure 9**). The most common canonical activation pathways (indicated by shades of orange), shared among 4 cell clusters, included “Apelin cardiomyocyte signaling pathway” (clusters 1,3,5,and 7); “hepatic fibrosis signaling pathway” (activating in clusters 0, 1, 3, 7 and inhibiting in cluster 8); “production of nitric oxide and reactive oxygen species in macrophages” and “role of PKR in interferon induction and antiviral response” (clusters 0, 1, 3, and 8). Common inhibitory canonical pathways included “RhoGDI signaling” (clusters 1, 3 and 7); and “HOTAIR regulatory pathway” (clusters 1, 3, 5, and 8).

**Figure 9.**
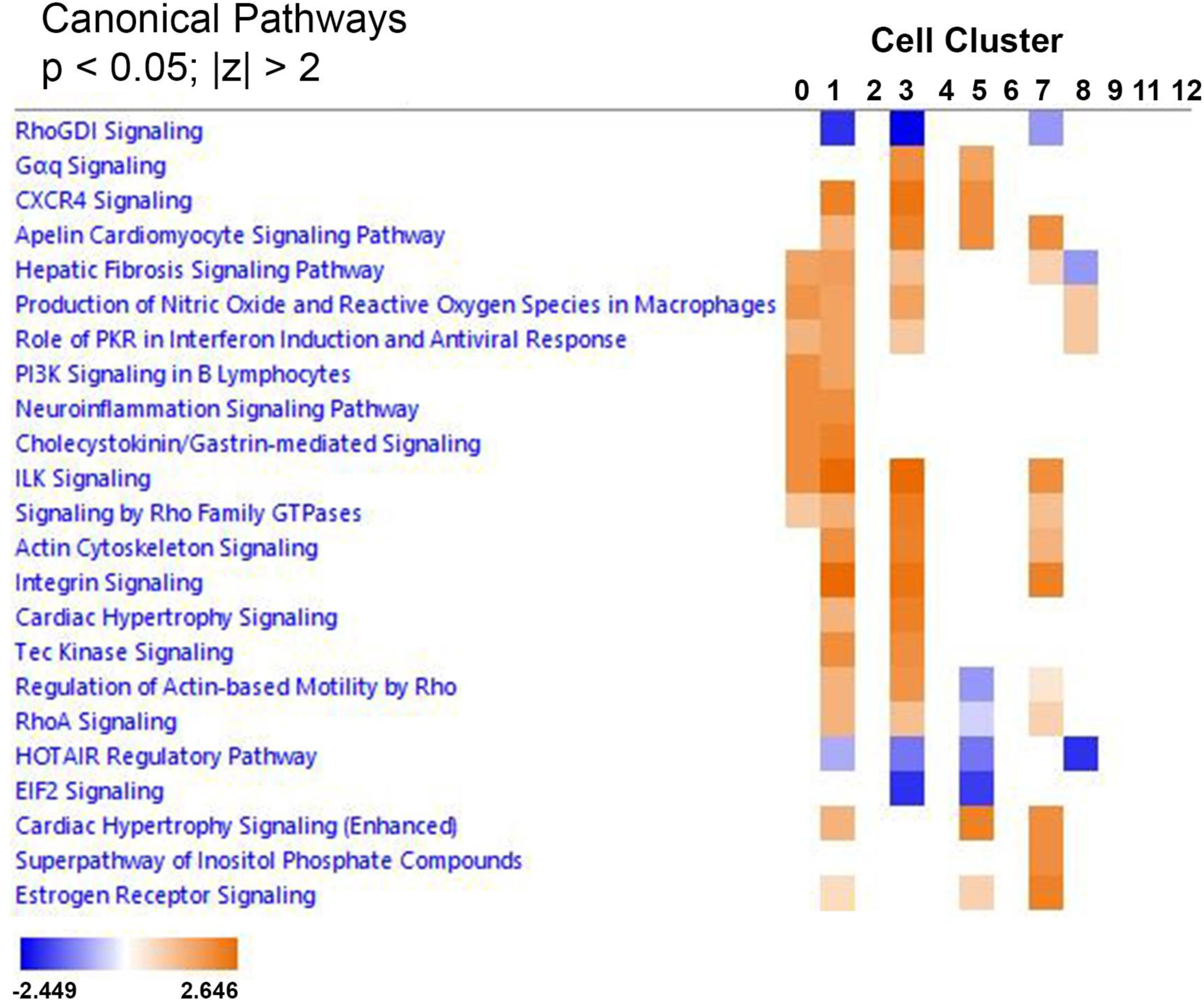
IPA comparison analysis predicted canonical pathways involved in the choroidal recovery response. Heat map of the comparison analysis of canonical pathways significantly enriched in the indicated cell clusters from recovering eyes compared with controls. Enrichment values [activation z-score] are scaled from −2.449 to 2.646 (blue to orange) and grouped by hierarchical clustering. A positive z-score (orange) denotes pathway activation, and a negative z- score (blue) denotes pathway inhibition.

A comparison analysis of upstream regulators was performed for the 12 cell clusters in order to predict which regulators might mediate the gene expression changes observed in all choroidal cell clusters. From **Figure 10** it can be predicted that top activators of the choroidal recovery response are INFG, TNF, Il1B, EGF, HGF and OSM (orange indicates activation), while MYC, MLXIPL, MYCN, SOCS3, DNMT3A, alpha catenin, miR-1-3P, and ERBB4 are predicted to inhibit the choroidal recovery response (blue indicates inhibition).

**Figure 10.**
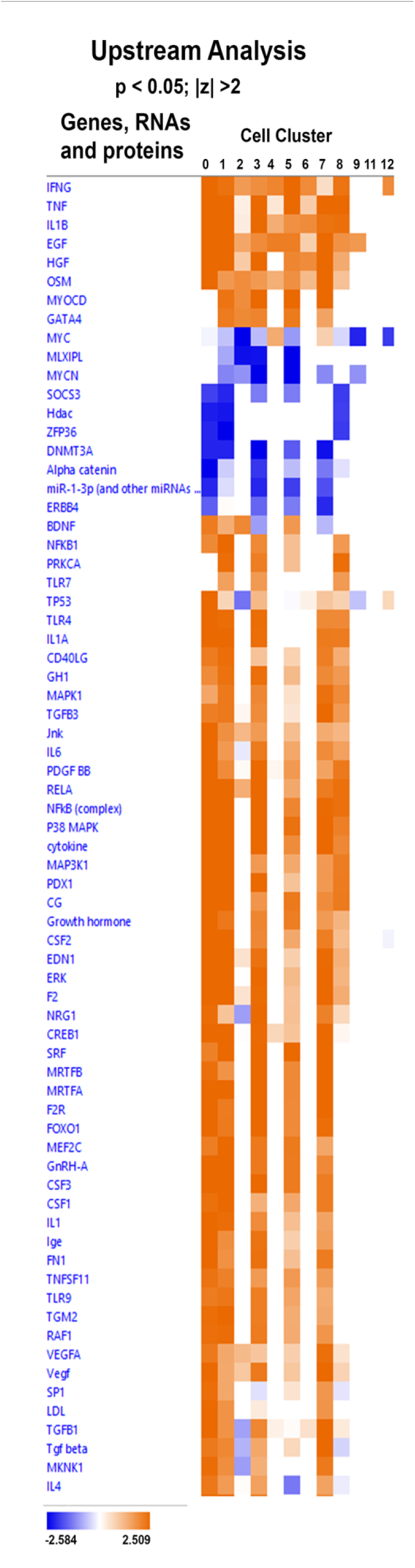
IPA comparison analysis predicted upstream regulators involved in the choroidal recovery response. Heat map of the comparison analysis of the statistically significant upstream regulators predicted to be involved in the indicated cell clusters. Enrichment values [activation z-score] are scaled from ─2.584 – 2.509 (blue to orange, and grouped by hierarchical clustering.

## Discussion

In an effort to elucidate the molecular mechanisms underlying the process by which vertebrate eyes can modulate their postnatal growth to maintain emmetropia (emmetropization), we utilized single cell RNA sequencing to identify the cell types present in the chick choroid and compare the transcriptomic changes in 24 choroidal cell populations between control eyes and eyes in which their growth is slowed as a consequence of form deprivation myopia followed by a brief period of unrestricted vision (recovery).

We observed significant increases in three subpopulations of vascular endothelial cells in recovering eyes as compared with control eyes following 24 hrs of recovery, suggesting that recovery is associated with choroidal angiogenesis. This observation is in agreement with our previous observation of an increased number of proliferating endothelial cells in recovering choroids based on BRDU incorporation (Summers, Cano et al., 2020). Additionally, a population of non-myelinating Schwan cells (NMSCs) in cell cycle phase G1/S (cluster 6) was significantly decreased in recovering eyes, compared with controls. Non-myelinating Schwann cells typically surround several small diameter axons, ensheathing each axon in a pocket of cytoplasm, forming a Remak bundle. These cells provide support and nutrition to axons, ensuring their survival. NMSC’s have been described in close association with blood vessels as well as in a “synapse-like” interaction/association of NMSC processes with various subsets of dendritic cells, macrophages, and lymphocytes (Ma et al., 2018). Interestingly, nonmyelin-forming Schwann cells, have the capacity to act as the “first responders” to injury or disease (Griffin and Thompson, 2008). Our observed changes in NMSC cell number and gene expression following only 24 hrs of recovery suggest that these cells may be acting as “first responders” in the choroid.

Taken together, our results indicate that our visual manipulations significantly affected gene expression in nearly all endogenous cell populations in the choroid. Only a subpopulation of melanocytes and a subpopulation of non-myelinating Schwann cells (NMSC), both in cell cycle phases G2/M, were unaffected. No changes in gene expression were detected in 5 of the 6 blood cell types identified in the choroid. Only the population of choroidal M1 macrophages (M1 Mɸ) displayed significant differences in gene expression as well as a significant increase in cell number in response to recovery from induced myopia. However, due to the large differences in cell numbers within each choroid cell population (e.g. population 0 is the largest population with 513 – 949 cells and population 23 is the smallest with 2 – 20 cells), statistical significance could be reached with smaller differences in gene expression among clusters with larger sample sizes and only genes with large fold changes could be reported as significantly different in clusters with small samples sizes. Nevertheless, our results suggest that recovery is associated with relatively small fold changes (less than 2 fold) in most choroidal cell populations. Interestingly, the largest fold changes were detected in cells in cluster 21 (neurons/neuroendocrine cells; 7 – 12 fold), suggesting that these cells respond robustly to visual and/or chemical signals produced during the emmetropization process. We confirmed that cluster 21 neurons represent a rare cell population in the choroid, by using anti-CHRNA3 antibodies to immunolocalize these cells in choroidal whole mounts and to isolate these cells from choroidal cell suspensions via FACS. In addition to the expression of multiple neuron-specific genes (CHRNA3, GAL, NPY, SCN9A and STMN2), cluster 21 cells highly overexpressed three opsin genes, OPN1LW (long wave sensitive opsin), OPN1MSW (green-sensitive, rhodopsin-like opsin), and OPN4-1 (a retrogene with similar absorption spectrum to melanopsin) compared with all other cell types in the choroid, suggesting that cluster 21 neurons may be light sensitive, particularly to long and medium-wavelength light (yellow - green range). Several studies have documented changes in choroidal thickness in humans in response to blue or blue green light (peak wavelengths of 465 and 500 nm, respectively) (Lou and Ostrin, 2020; Read et al., 2018), which are close to the peak sensitivity of melanopsin (Opn4; 480 nm) (Berkowitz et al., 2016; Berson et al., 2002; Dacey et al., 2005). This light evoked choroidal thickening may represent a downstream response, initiated in the rods and cones, or in the melanopsin-containing intrinsically photosensitive retinal ganglion cells (ipRGCs). However, based on our identification of opsin-gene expression in choroidal neurons, it is possible that the choroid has some intrinsic photosensitivity as well, which could play a role in the observed light-driven changes in choroid thickness.

In an attempt to identify global patterns of gene expression changes in the choroid associated with recovery from induced myopia, we compared all genes that displayed significant differences among three or more choroidal cell clusters (**Figure 6**). Heat map analyses indicated that, with a few exceptions, gene expression differences were similar between different choroidal cell populations (i.e., genes that were upregulated in one cluster were similarly upregulated in several cell clusters and genes that were downregulated were downregulated in many cell clusters). This is not surprising, considering that many of the large cell populations represented subpopulations of the same cell type. Genes that were significantly overexpressed in six or more choroid cell clusters included MT4, MYH11, IRF1, ATF3 and EGR1:

### MT4 (metallothionein 4)

MT4 is a low molecular weight cysteine rich metalloprotein. In mammals, there are four isoforms (MT-1, -2, -3, and -4) and they have multiple roles, such as the detoxification of heavy metals, regulating essential metal homeostasis, and protecting against oxidative stress. Accumulating studies have suggested that metallothioneins are an important neuroprotective substance for cerebral ischemia and retinal diseases, such as age-related macular degeneration (AMD) and retinitis pigmentosa (RP), that are characterized by a progressive retinal degeneration (Ito et al., 2013).

### MYH11 (Myosin-11)

MYH11 encodes the smooth muscle cell-specific myosin heavy chain, a major component of the contractile unit of smooth muscle cells (Babu et al., 2000). The myosin light chain kinase MYLK was also upregulated in five choroid cell clusters (0,1,3,5,7). The upregulation of these smooth muscle myosin proteins, together with the observation of increased numbers of vascular endothelial cells in recovering choroids, provide further support that recovery from induced myopia is associated with angiogenesis in the choroid.

### IRF1 (interferon regulatory factor 1)

IRF1 is an interferon (IFN) and virus inducible gene, also induced by tumor necrosis factor (TNF), interleukin 1 (IL-1), poly(I:C) (Fujita et al., 1989) and retinoic acid (Kroger et al., 2002). IRF1 is preferably induced by type I IFNs (IFN-I) and orchestrates the proinflammatory response mediated by IFN-I (Kroger et al., 2002) through the upregulation of IFNβ, iNOS, and IL-12p35 (Negishi et al., 2006). IRF1 has been shown to have roles in apoptosis, inflammation, cell growth and polarization, oncogenesis, and cancers (Alsamman and El-Masry, 2018; Savitsky et al., 2010; Tamura et al., 2008, Yan et al., 2021; Yanai et al., 2012).

### ATF3 (Activating transcription factor 3)

ATF3 is a stress-induced transcription factor that plays vital roles in modulating metabolism, immunity, and oncogenesis (Chen et al., 1996). ATF3 acts as a hub of the cellular adaptive-response network and is induced by a variety of extracellular signals, such as endoplasmic reticulum (ER) stress, cytokines, chemokines, and LPS (Ku and Cheng, 2020). In the eye, ATF3 expression has been shown to be upregulated in the ganglion cell layer of the retina and the optic nerve of eyes following optic nerve injury, possibly providing a neuroprotective role in the retina following injury and a mediator of optic nerve regeneration (Kole et al., 2020; Saul et al., 2010).

### EGR1 (early growth response protein 1)

We observed EGR1 upregulated in 8 cell clusters in response to recovery from induced myopia, representing the most globally upregulated gene in the recovering choroid. EGR1 [also known as “ZENK”, and “nerve growth factor-induced protein A (NGFI-A)”] is known as an immediate early gene (IEG). IEGs are known mediators of gene and environment interactions; providing the ability to trigger a fast response involving many downstream effects. As a transcriptional regulator, EGR1 has been shown to be involved in a wide variety of processes including neural plasticity and neuronal activity, extracellular matrix remodeling and fibrosis, host-pathogen interactions and carcinogenesis (Herdegen and Leah, 1998; Bahrami and Drablos, 2016; Banerji and Saroj, 2021; Havis and Duprez, 2020; Wang et al., 2021). Egr-1 gene expression has been shown to exhibit a bi-directional response to opposing ocular growth stimuli in retinas of both chick and guinea pig models of myopia and emmetropization; EGR1 mRNA is downregulated in a population of retinal amacrine cells under visual conditions associated with increased ocular growth and myopia development, and is upregulated in the retinal amacrine cells in response to visual conditions associated with decreased ocular growth and recovery from myopia (Ashby et al., 2014; Fischer et al., 1999). Our finding that EGR1 is upregulated in several populations of choroidal endothelial cells (clusters 0 and 4), fibroblasts (clusters 1,3,5, and 7), mural cells (cluster 8), and non-myelinating Schwann cells (cluster 6) in recovering eyes, suggests that EGR1 transcriptional activity may coordinate the growth of multiple ocular tissues in response to visual signals to effect changes in refraction.

### Pathway Analyses

Comparison analyses were used to identify pathways and biological functions common across multiple cell types involved in the choroidal recovery response from myopia. Most notably, the canonical pathway, “production of nitric oxide and reactive oxygen species in macrophages” and was predicted to be activated based on the gene expression changes among lymphatic vascular endothelial cells (cluster 0), canonical fibroblasts (clusters 1 and 3), and mural cells (cluster 8). Nitric oxide has previously been implicated in mediating recovery from myopia since intravitreal administration of the non-specific nitric oxide synthase inhibitor, L-NAME, temporarily inhibits the choroidal and scleral responses associated with recovery from induced myopia (Nickla et al., 2006; Summers and Martinez, 2021) and intravitreal delivery of the NOS substrate, L-arginine or a nitric oxide donor (sodium nitroprusside; SNP) significantly inhibits form deprivation myopia (Carr and Stell, 2016). Additionally, nitric oxide has been shown to mediate the ocular growth-inhibiting properties of both the muscarinic receptor antagonist, atropine (Carr and Stell, 2016) and the dopamine agonist, quinpirole (Nickla et al., 2013). Therefore, we predict that nitric oxide is one of the earliest signaling events in the process of emmetropization since administration of L-NAME immediately prior to recovery blocks the recovery-induced increase in IL6 observed following 6 hrs of recovery (Summers and Martinez, 2021) as well as the change in choroidal thickening and axial elongation observed 7 hrs after recovery (Nickla and Wildsoet, 2004). Data from the present study also indicate that the number of M1 macrophages is increased in recovering choroids. Since macrophages are a known source of nitric oxide via expression of inducible nitric oxide synthase (iNOS), it is possible that choroidal macrophages could be a source of nitric oxide, in addition to photoreceptors and the RPE.

Comparison analyses were also used to identify potential upstream regulators that could be predicted to induce the gene expression changes in multiple choroid cell clusters associated with recovery from induced myopia. This analysis indicated that INFG, TNF, IL1B, EGF, HGF and OSM would be predicted to stimulate recovery from induced myopia (activators). It is generally accepted that visually guided eye growth is regulated by a cascade of chemical events that is initiated in the retina, is transmitted and modified through the RPE and choroid, and ultimately acts on the sclera to affect changes in eye size and refraction. Therefore, these predicted upstream activators may be synthesized in the retina and/or RPE and secreted into the choroid to stimulate gene expression changes in multiple choroid cell populations during the choroidal recovery response.

## Conclusions

Previously, we observed rapid and significant changes in choroidal gene and protein expression of the pro-inflammatory cytokines IL6 and IL1β in response to myopic defocus, either during recovery from myopia or following application of +15D lenses to normal chick eyes (Summers and Martinez, 2021). Interestingly, systemic treatment of chicks with the potent anti-inflammatory agent, dexamethasone, significantly reduced choroidal Il6 gene expression in recovering eyes and inhibited the scleral extracellular matrix changes associated with recovery from myopia (Summers et al., 2022). These results suggest that the regulation of postnatal eye growth may involve aspects of the innate immune system acting in coordination with the visual environment to regulate scleral remodeling, ocular size and refraction. In the present study, our analyses of the recovery-induced differentially expressed genes, canonical pathways and upstream regulators of the choroid gene expression changes identified in multiple cell populations also suggest that recovery/compensation for myopic defocus may resemble a modified/limited inflammatory response involving multiple cell types (fibroblasts, mural cells, endothelial cells, macrophages), nitric oxide, and cytokines (IFNG, TNF, IL1B, EGF). Further studies to identify the key upstream mediators responsible for orchestrating the choroidal response when the eye is slowing its rate of elongation will not only help to elucidate the mechanism of emmetropization, but provide new therapeutic targets for the treatment of myopia in children.

## Materials and methods

### Key Resources Table

| Key Resources Table |  |  |  |  |
| --- | --- | --- | --- | --- |
| Reagent type (species) or resource | Designation | Source or reference | Identifiers | Additional information |
| antibody | anti-nicotinic acetylcholine receptor, $\alpha 3$ subunit (CHRNA3; Rat polyclonal, primary antibody) | Sigma | Cat# M221 088H4612 | IF and FACS (1:4000) |
| antibody | Alexa Fluor 568 – Goat anti-rat IgG (secondary antibody) | ThermoFisher Scientific | Cat# A-11077 | IF (1:1000) |
| antibody | Alexa Fluor 647– Goat anti-rat IgG (secondary antibody) | ThermoFisher Scientific | Cat# A-21247 | FACS (1:1000) |
| chemical compound | Collagenase Type 1 | Worthington Biochemical Corporation | Cat# LS004196 Lot# 48S18874 | 2 mg/ml |
| chemical compound | Trypsin (porcine pancreas) | Millipore Sigma | Cat# T6567 | 0.20 % |
| other | HBSS (10X) calcium, magnesium, phenol red | ThermoFisher Scientific | Cat# 14060040 | 1X for tissue digestion |
| other | DAPI stain | Invitrogen | Cat#: D3571 | IF and FACS (1:1000) |

### Ethics and Animals

Animals were managed in accordance with the ARVO Statement for the Use of Animals in Ophthalmic and Vision Research, with the Animal Welfare Act, and with the National Institutes of Health Guidelines. All procedures were approved by the Institutional Animal Care and Use Committee of the University of Oklahoma Health Sciences Center (protocol # 20-092-H). White Leghorn male chicks (*Gallus gallus*) were obtained as 2-day-old hatchlings from Ideal Breeding Poultry Farms (Cameron, TX). Chicks were housed in temperature-controlled brooders with a 12-hour light/dark cycle and were given food and water ad libitum. At the end of experiments, chicks were euthanized by overdose of isoflurane inhalant anesthetic (IsoThesia; Vetus Animal Health, Rockville Center, NY), followed by decapitation.

### Visual Manipulations

Form deprivation myopia (FDM) was induced in 3 to 4 day-old chicks by applying translucent plastic goggles to the right eye, as previously described (Rada et al., 1991). The contralateral eyes (left eyes) of all chicks remained untreated and served as controls. Chicks were checked daily for the condition of the goggles. Goggles remained in place for 10 days. At the start of the light cycle on the 11^th^ day (7 am) the goggles were removed and chicks were allowed to experience unrestricted vision (recover) for 24 hrs.

### Choroid Cell Isolation

Following 24 hrs of unrestricted vision, chicks were euthanized by an overdose of isoflurane inhalant anesthetic (IsoThesia; Vetus Animal Health). Eyes were enucleated and cut along the equator to separate the anterior segment and posterior eye cup. Anterior tissues were discarded, and the vitreous body was removed from the posterior eye cups. An 8 mm punch was taken from the posterior pole of the chick eye using a dermal biopsy punch (Miltex Inc., York, PA). Punches were located nasal to the exit of the optic nerve, with care to exclude the optic nerve and pecten oculi. With the aid of a dissecting microscope, the retina and majority of RPE were removed from the underlying choroid and sclera with a drop of phosphate buffered saline (PBS; 3 mM dibasic sodium phosphate, 1.5 mM monobasic sodium phosphate, 150 mM NaCl, pH 7.2) with gentle brushing. Choroids were separated from the sclera using a small spatula, placed in 4 ml snap cap tubes containing 2 ml of HBSS, and placed on ice. To ensure that there were a sufficient number of cells for RNA seq analysis, and to generate enough biological replicates for statistical analyses, choroids from control and recovering eyes of ten chicks each were pooled separately to generate three pools of control eyes (10 control choroids/pool) and three pools of recovering eyes (10 recovering choroids/pool; 30 chickens total).

Choroidal cells were isolated as described previously (Coulombe and Nishi, 1991) with minor modifications: Choroids were cut into small pieces, and digested with collagenase [Worthington; 2mg/ml in HBSS containing Ca^++^ and Mg^++^; 2ml/ sample] for 20 min at 37°C. Following centrifugation at 834 x g at 4°C for 15 min, the supernatants were removed and samples were digested with 2 ml trypsin (Sigma; 0.2% in PBS for 20 min at 37°C). Choroids were again centrifuged as described above, trypsin removed, and replaced with another 2ml of collagenase solution (2 mg/ml in HBSS) and incubated at 37°C with occasional trituration using a glass Pasteur pipette until the digest was able to pass freely through the glass Pasteur pipette. Following a second centrifugation as described above, supernatants were removed and the pelleted choroidal digests were resuspended in PBS + 0.04% BSA (1 ml/sample). The dissociated cell suspensions were passed through small, individual 105 μm metal mesh screens held in place with 25 mm Swinnex filter holders (MilliporeSigma, Burlington, MA) using 5 cc syringes (6 filters and 6 syringes in total). Each filtrate was collected in 4 ml snap cap tubes. Each filter was rinsed with an additional 1 ml PBS + 0.04% BSA and added to each original filtrate. Each cell sample was then passed through a 40 μm pipette tip cell strainer (FLOWMI; SP Bel-Art, Wayne, NJ). Living and dead cells were identified by simultaneously staining with green-fluorescent calcein-AM and red-fluorescent ethidium homodimer-1, respectively. For live/dead staining, cells were pelleted by centrifugation at 350 x g for 10 min at 4 °C, supernatants were removed, and cell pellets were gently resuspended in 0.5 ml PBS+ 0.04% BSA containing live/dead stain ((LIVE/DEAD™ Viability/Cytotoxicity Kit, Thermofisher, Waltham, MA; 5μl calcein AM + 20 ul EthDIII in 10 ml PBS+ 0.04% BSA). Cell samples were incubated in live/dead stain for 45 min at room temperature immediately prior to fluorescence activated cell sorting (FACS) (see below).

### FACS for Single-Cell Experiments

Following staining of cells with live/dead stain, living (green fluorescent) cells were separated from dead (red fluorescent) cells and isolated by FACS on a BD FACSAria™ Fusion (BD Biosciences Franklin Lakes, NJ) using an excitation with a 488 nm laser and the emission collected using a 530/30 band-pass filter for living (calcein AM-labelled) cells and excitation with a 561 nm laser and emission collected with a 610/20 band-pass filter for dead (EthDIII-labelled) cells, together with a 100 μm nozzle size. Single live cells were defined by electronic gating in FACS DIVA software (BD, ver. 8.01) using forward and side-angle light scatter (FSC and SSC, respectively), calcein AM (green) fluorescence and the absence of EthDIII (red) fluorescence. Calcein AM- positive cells were sorted directly into PBS+ 0.04% BSA.

Following FACS, cells were counted with an automated Cell Counter (BioRad TC20, Hercules, CA) and living cell concentrations were adjusted to 1000 cells/μl in a final volume of 50 μl) and transported on wet ice to the OUHSC Institutional Research Core Facility for single cell RNA sequencing and bioinformatic analyses.

### Single Cell RNA-seq libraries

Sorted cells were given to the Genomics division of the Institutional Research Core Facility. Single cell libraries were built using 10x Genomics’ Chromium Controller, Chromium Next GEM Single Cell 3’ Reagent Kits v3.1 and their established protocols. We captured 10,221 - 12,745 cells per sample using the 10X Chromium single cell system and sequenced each sample to a read depth of 35,621 - 46,031 reads/cell (**Table S1**).

### Bioinformatic Analysis of scRNA-seq Data

Read mapping, and expression quantification were performed using a combination of the 10X Cellranger pipeline and custom Seurat analytic scripts. Briefly, single-cell reads were mapped to the chicken genome (GRCg6a) and assigned to genes using the standard CellRanger pipeline. Normalized gene expression was then exported to Seurat and subsequently used to produce a UMAP plot that provides cell clusters based on similarity of gene expression. Once cells were assigned to a cluster, custom Seurat scripts were used to statistically derive the gene expression differences within and between cell clusters using T-tests. Seurat was then used to generate a marker list for each cluster using the “FindAllMarkers” algorithm. The Seurat “Cell Cycle Scoring” routine was used to identify subclusters of cells in G1, S and G2/M phases. The gene look up databases, GeneCards, GenePaint website (https://gp3.mpg.de), and Human Protein Atlas (https://www.proteinatlas.org) were then consulted to determine the possible identity of a cluster. Additionally, StringDB (https://string-db.org) was used to look up marker genes in order to identify possible interacting genes, and then determine if those interacting genes were also expressed in the same cluster. Additional to cluster identification, pathway analysis was performed using Ingenuity Pathway Analysis (Qiagen, Redwood City, CA) on the differential transcriptional profiles seen in the cell clusters.

### Choroidal Immunolabelling

Punches (8 mm) were taken from the posterior poles of control and recovering chick eyes and cleaned of retina and RPE as described above. The sclera, with choroid still attached, was placed in 4% paraformaldehyde at 4 °C overnight. After fixation, chick choroids were gently removed from the scleral tissue, and placed into a 48-well flat bottom plate (Corning Inc., Corning, NY). Choroids were washed in PBS for 10 min (3X) on a shaking device at RT. Choroids were then blocked in BSA-PBS (2% BSA, 0.2% Triton X-100, 0.004% sodium azide in PBS, pH 7.4) for 1 hr at RT with rocking and subsequently incubated in BSA-PBS containing primary antibodies (**Key Resources Table**) for 72 hrs at 4°C. Choroids were then washed in PBS for 10 min (6X) at RT with rocking, incubated with goat anti-rabbit IgG conjugated to AlexaFluor 568 (1:1000 in BSA-PBS; Life Technologies Grand Island, NY) for 24 hours at 4°C, and washed with PBS for 10 min (6X). Following the PBS washes, choroids were stained with DAPI (diluted 1:1000 in PBS from 5mg/ml stock in dH_2_0), mounted on glass slides and coverslipped using a fluorescence mounting media (Prolong Gold with DAPI, Thermo Fisher Scientific). All slides were stored at 4°C until imaging. Immunolabeled choroids were examined under an Olympus Fluoview 1000 laser-scanning confocal microscope (Center Valley, PA).

### FACS sorting of CHRNA3-positive cells

Choroidal cells were isolated as described above and fixed in 4% formaldehyde at 4°C overnight. Cells were then rinsed in wash buffer (PBS + 2% BSA), following by incubation in blocking buffer (2% BSA, 0.2% Triton X-100 in PBS) for 10 min at room temperature. Cells were immunolabelled with anti-chick CHRNA3 (1:4000) in blocking buffer overnight 4°C with rotation. Cells were then washed 3 x 10 min with wash buffer followed by incubation in goat anti-rat alexafluor 647, diluted 1:1000 in wash buffer for 1 hr at room temp with rotation. Cells were then washed 3 X 10 minutes with wash buffer, nuclei were labelled with DAPI (1:1000 in wash buffer), and stored in wash buffer at 4°C in the dark prior to FACS analysis. Stained cells were subjected to data acquisition using a Stratedigm S1400Exi flow cytometer (Stratedigm, Inc., San Jose, CA) using an excitation of 640 nm and the emission collected using a 676/29 nm band pass filter. Data were analyzed by FlowJo software.

## Acknowledgements

This work was supported by NIH grant R01EY09391 (JAS). The authors also thank the Institutional Research Core Facility at OUHSC for the use of the Core Facility which provided single cell RNA library construction.

