## Supplemental Figures and Tables for "Single Cell Transcriptomics Identifies Distinct Choroid Cell Populations Involved in Visually Guided Eye Growth"

**Supplementary Material**

**Figure 2- Figure supplement 1**

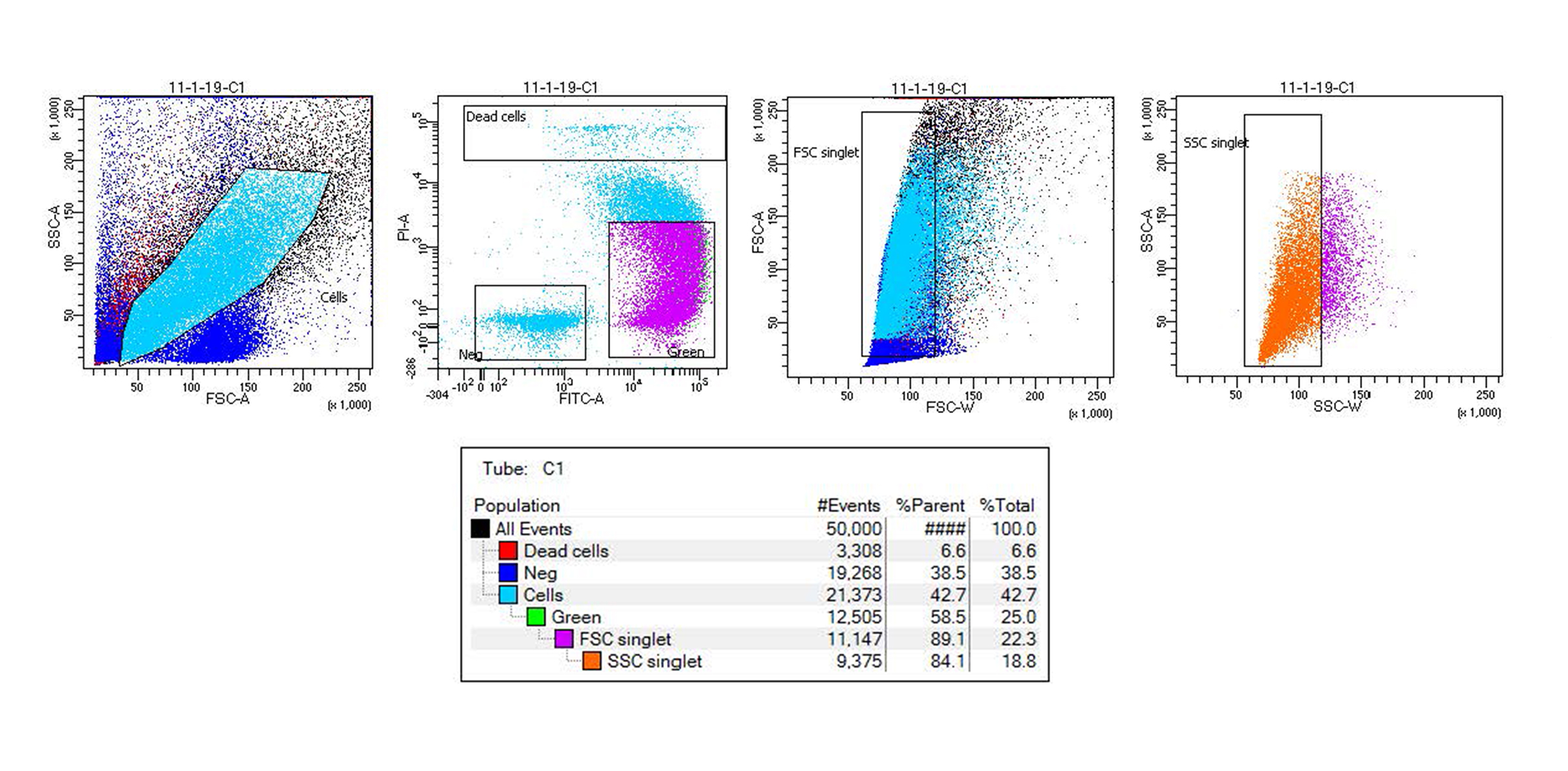

**Figure 2 - Figure supplement 1. FACS gating strategy for isolation of choroidal cells.**  FACS plot of all events (black) with forward scatter area intensity (FSC-A) and side scatter area intensity (SSC-A) containing a gate to select only single cells (royal blue) followed by a FACS plot based on green fluorescence intensity (FITC-A) and red fluorescence intensity (PI-A). Living cells labelled with calcein AM (green), but not with EthDIII (red; dead cells) were backgated onto gates of forward scatter width intensity (FSC-W) and forward scatter area intensity (FSC-A) followed by gates of side scatter width intensity (SSC-W) and sice scatter area intensity (SSC-A) to select only single living cells while avoiding cell aggregates.

**Table S1.** Sample data for Single Cell RNA sequencing.

| **Sample** | **# Cells/Sample** | **Mean Reads per Cell** | **Median Genes per Cell** |
| --- | --- | --- | --- |
| C1 | 10,221 | 46,031 | 1,110 |
| C2 | 11,251 | 43,351 | 1,095 |
| C3 | 12,469 | 41,847 | 1,225 |
| R1 | 11,976 | 35,621 | 1,080 |
| R2 | 12,392 | 40,631 | 1,140 |
| R3 | 12,745 | 43,480 | 1,123 |

**Table 1 – Figure supplement 1.** Dot blot showing the expression pattern and level of some established marker genes (shown in columns) for each major cell type (shown in rows) in chicken choroid. Color intensities show the expression level of the indicated gene.

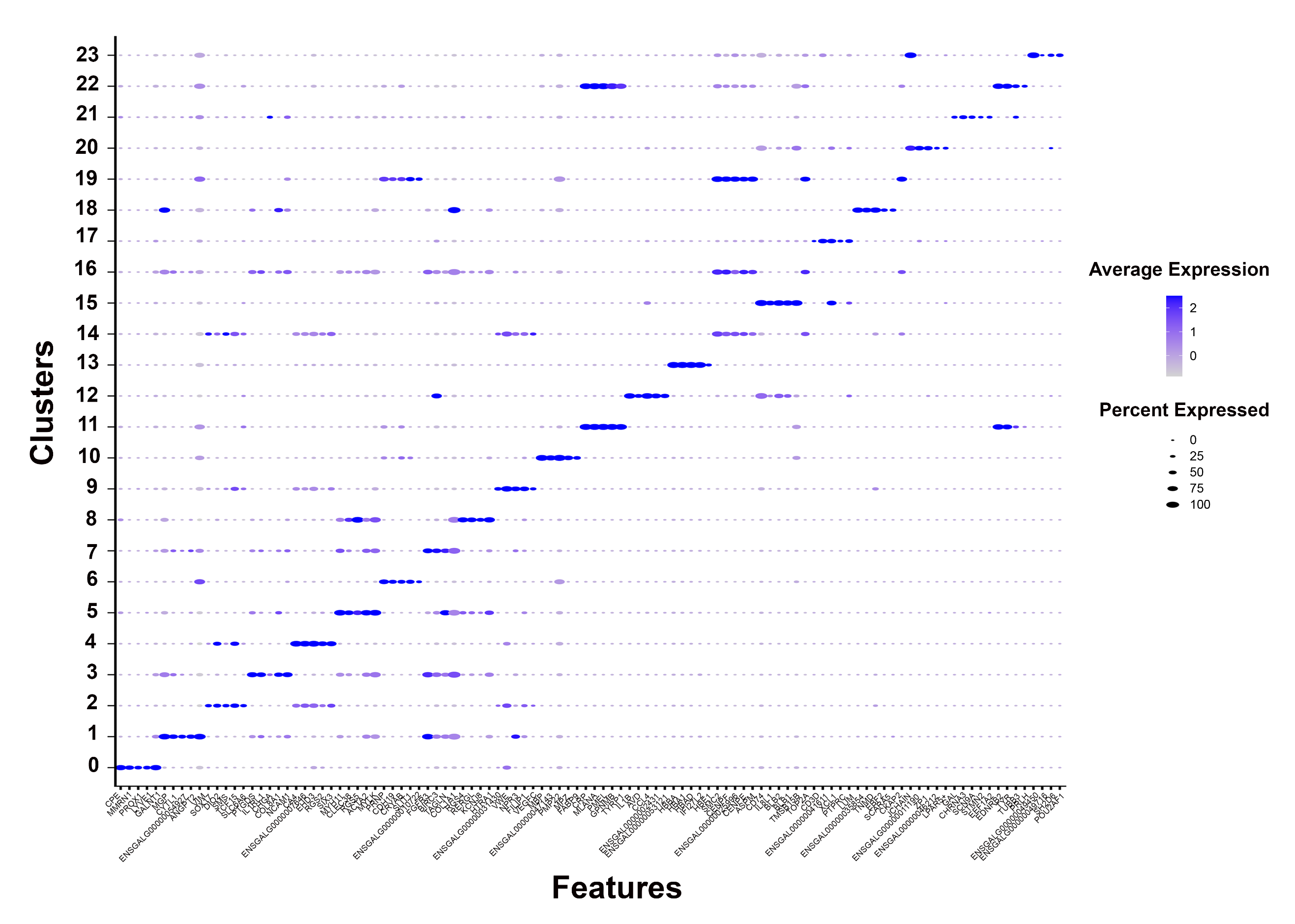

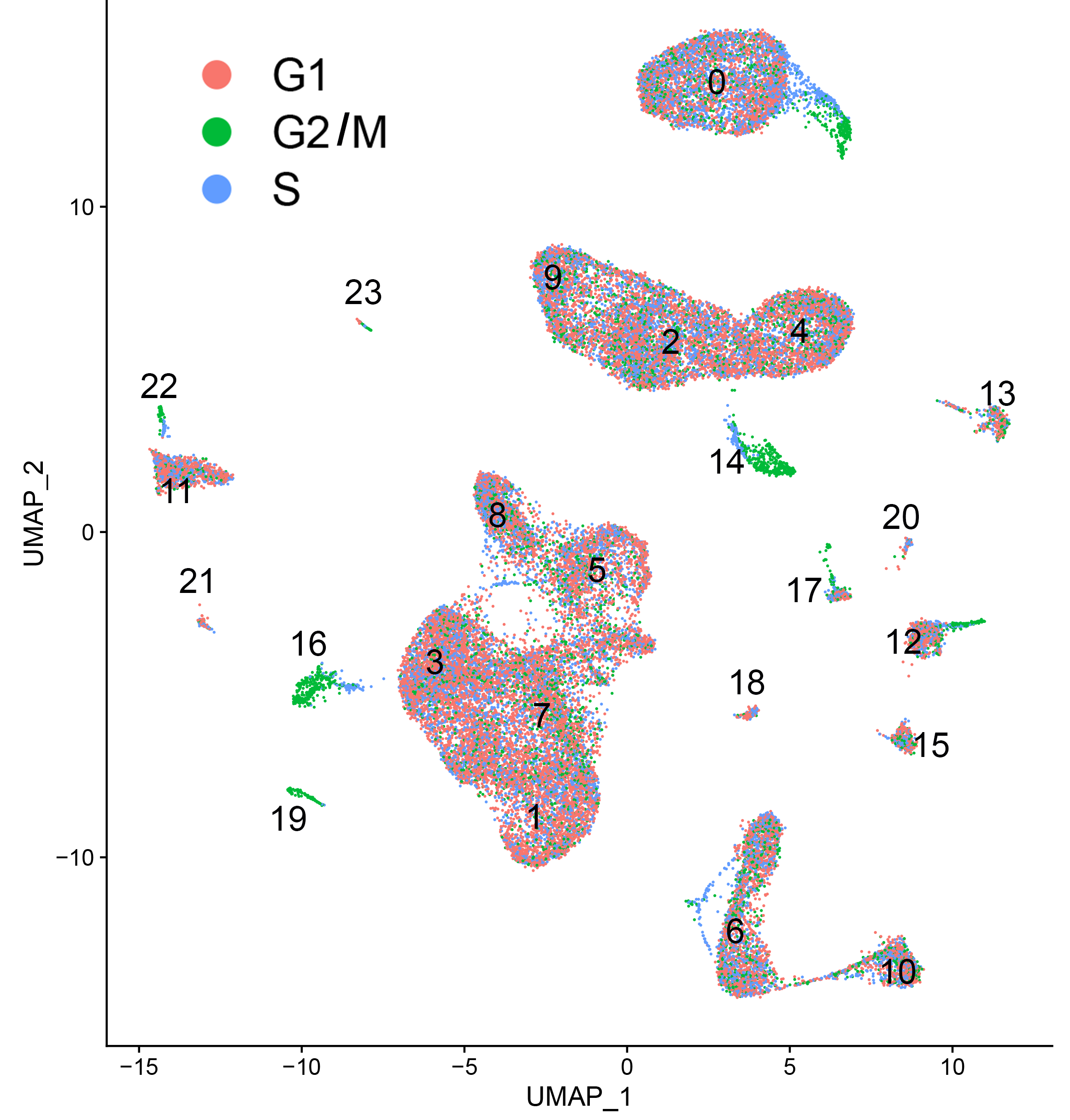

**Figure 3 – Figure supplement 1**. Uniform Manifold Approximation and Projection (UMAP) maps showing phases of cell cycle assigned for each of the 24 choroid cell clusters as determined from the “Cell Cycle Scoring” function in Seurat v 4.0.6). G1, growth 1 phase; G2/M, growth 2/ mitosis phase; S, synthesis phase.

**Figure 3 – Figure supplement 2.** Uniform Manifold Approximation and Projection (UMAP) maps showing expression of fibroblast marker genes in choroidal cell populations. Col1A1, collagen type I alpha 1 chain; DES, desmin; RGS4, regulator of G protein signaling 4; RGS5, regulator of G protein signaling 5; LMOD1, leiomodin 1; LUM, lumican.

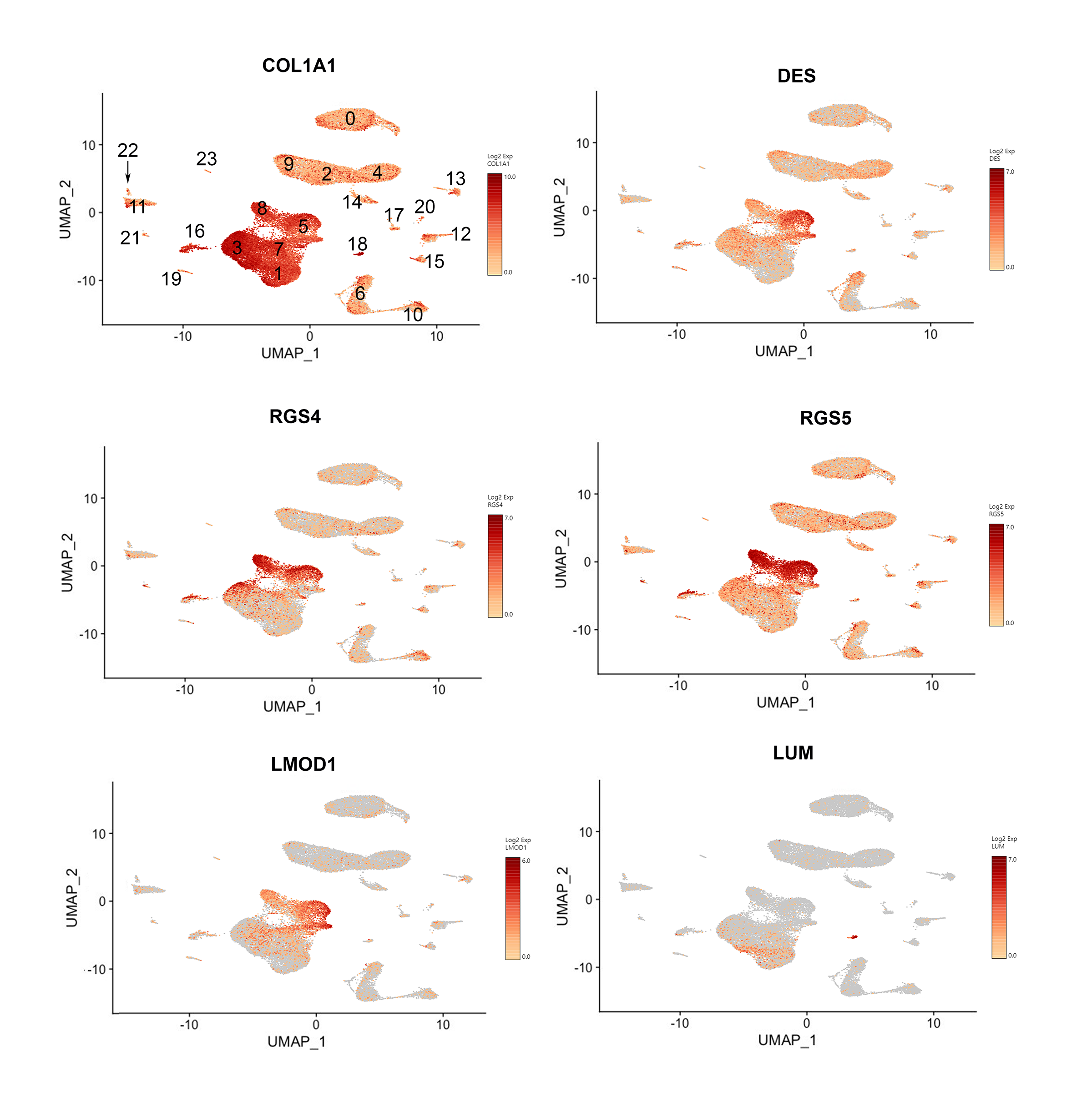

**Table S2**. Percentages of cells in each cluster of total choroidal cell populations.

| **seurat_clusters** | **CONT_C1** | **CONT_C2** | **CONT_C3** | **RECOV_R1** | **RECOV_R2** | **RECOV_R3** | **Control Mean** | **RECOV Mean** |
| --- | --- | --- | --- | --- | --- | --- | --- | --- |
| **0** | 9.41% | 10.97% | 11.51% | 14.46% | 13.56% | 14.31% | 10.63% | 14.11% |
| **1** | 12.56% | 11.75% | 11.78% | 10.97% | 12.74% | 10.18% | 12.03% | 11.30% |
| **2** | 9.45% | 10.83% | 10.38% | 12.23% | 10.97% | 14.49% | 10.22% | 12.56% |
| **3** | 11.68% | 9.83% | 11.58% | 9.96% | 10.82% | 9.54% | 11.03% | 10.11% |
| **4** | 7.34% | 7.28% | 8.21% | 9.26% | 7.11% | 9.73% | 7.61% | 8.70% |
| **5** | 8.24% | 8.17% | 10.81% | 6.98% | 6.87% | 5.94% | 9.07% | 6.60% |
| **6** | 8.79% | 9.36% | 7.50% | 6.96% | 7.44% | 5.27% | 8.55% | 6.56% |
| **7** | 7.54% | 7.70% | 6.74% | 3.93% | 6.74% | 4.53% | 7.33% | 5.07% |
| **8** | 5.85% | 4.48% | 5.20% | 4.97% | 4.08% | 3.22% | 5.18% | 4.09% |
| **9** | 3.26% | 2.93% | 2.86% | 5.00% | 4.78% | 6.11% | 3.02% | 5.29% |
| **10** | 3.82% | 4.27% | 2.95% | 2.41% | 2.55% | 2.35% | 3.68% | 2.43% |
| **11** | 3.15% | 3.50% | 2.52% | 2.74% | 2.45% | 3.35% | 3.06% | 2.85% |
| **12** | 1.98% | 1.36% | 1.67% | 2.77% | 2.62% | 2.30% | 1.67% | 2.56% |
| **13** | 1.56% | 2.06% | 1.11% | 1.01% | 0.86% | 1.43% | 1.57% | 1.10% |
| **14** | 0.75% | 0.68% | 1.00% | 1.39% | 1.38% | 2.18% | 0.81% | 1.65% |
| **15** | 1.39% | 0.92% | 0.98% | 1.07% | 1.16% | 1.20% | 1.10% | 1.14% |
| **16** | 1.14% | 1.02% | 0.83% | 1.40% | 1.24% | 0.94% | 1.00% | 1.19% |
| **17** | 0.61% | 0.55% | 0.64% | 0.87% | 0.84% | 1.53% | 0.60% | 1.08% |
| **18** | 0.44% | 1.00% | 0.53% | 0.43% | 0.48% | 0.34% | 0.66% | 0.42% |
| **19** | 0.24% | 0.34% | 0.33% | 0.37% | 0.36% | 0.30% | 0.30% | 0.34% |
| **20** | 0.29% | 0.34% | 0.19% | 0.27% | 0.29% | 0.33% | 0.28% | 0.30% |
| **21** | 0.22% | 0.44% | 0.48% | 0.11% | 0.22% | 0.11% | 0.38% | 0.15% |
| **22** | 0.20% | 0.18% | 0.14% | 0.27% | 0.15% | 0.21% | 0.17% | 0.21% |
| **23** | 0.09% | 0.03% | 0.07% | 0.18% | 0.29% | 0.13% | 0.06% | 0.20% |
| **Total %** | 100.00% | 100.00% | 100.00% | 100.00% | 100.00% | 100.00% |  |  |
